# Multilevel Regulation of Skeletal Muscle Ferroptosis in Aging: Sex- and Exercise-Dependent Effects on Histological, Molecular, and Genetic Markers

**DOI:** 10.64898/2025.12.08.692947

**Authors:** Fujue Ji, Hyeonseung Rheem, Haesung Lee, Minyeong Eom, Jong-Hee Kim

**Affiliations:** Department of Physical Education, College of Performing Arts and Sport, Hanyang University, 222 Wangsimni-ro, Seongdong-gu, Seoul, Republic of Korea; BK21 FOUR Human-Tech Convergence Program, Hanyang University, 222 Wangsimni-ro, Seongdong-gu, Seoul 04763, Republic of Korea; Aging and Aged Laboratory, Hanyang University, 222 Wangsimni-ro, Seongdong-gu, Seoul, Republic of Korea

**Keywords:** Ferroptosis, Aging, Sex, Exercise, Skeletal muscle

## Abstract

**Background:** Ferroptosis, an iron-dependent form of regulated cell death, is increasingly recognized as a key contributor to aging-associated skeletal muscle degeneration and dysfunction. However, the interactive effects of aging, sex, and exercise modality on ferroptosis regulatory markers at the histological, protein, and gene expression levels remain poorly understood.

**Methods:** Male (n = 23) and female (n = 23) mice aged 7 (young) and 17 (aged) months were assigned to sedentary control, voluntary wheel running, or forced treadmill exercise. Ferroptosis in the quadriceps muscle was assessed using histological markers (e.g., fibrosis, Fe³⁺ accumulation, 4-HNE, MDA), protein-level markers (e.g., GPX4, SLC7A11, p-AMPK, MDA, GSH/GSSG), and gene expression markers (e.g., SLC7A11, GSS, ACSL4, POR).

**Results:** Aging significantly elevated histological indicators of ferroptosis—fibrosis, lipid peroxidation, and iron overload—regardless of sex. At the protein and gene levels, sex-dependent differences were evident: aged females exhibited lower MDA and GSSG levels and upregulation of antioxidant-related genes, compared with aged males. Both exercise interventions modulated ferroptosis markers, with forced exercise exerting more pronounced effects than voluntary exercise. Notably, aged females demonstrated the most substantial reductions in ferroptosis-related markers in response to forced exercise, indicating a significant sex-by-exercise interaction.

**Conclusion:** Aging markedly increases ferroptosis-related changes in skeletal muscle, with partial sex-specific differences at the molecular level. Forced exercise provides more robust regulatory effect against ferroptosis than voluntary exercise, especially in aged females. These findings underscore the therapeutic potential of sex-specific, targeted exercise interventions for mitigating ferroptosis-mediated muscle deterioration during aging.

## Introduction

Ferroptosis is a distinct form of regulated cell death driven by iron-dependent lipid peroxidation and characterized by the accumulation of reactive oxygen species and oxidative degradation of polyunsaturated fatty acids in cellular membranes, ultimately leading to membrane rupture and cell demise [1–3]. Unlike apoptosis, necrosis, or autophagy, ferroptosis involves a unique metabolic and redox-dependent pathway that plays a pivotal role in various physiological and pathological conditions, including neurodegenerative diseases, cancer progression, and tissue injury responses [4–6].

Recent research has highlighted that ferroptosis is not solely a cell-autonomous event but is highly influenced by systemic physiological and biological factors [7, 8]. Aging, for instance, is associated with diminished antioxidant defense, dysregulated iron metabolism, and elevated lipid peroxidation—hallmark features that collectively contribute to ferroptosis susceptibility [9–11]. Similarly, sex-based biological differences—particularly in hormone levels, mitochondrial function, and oxidative stress responses—have been associated with differential regulation of ferroptosis-related genes and proteins [12, 13]. Physical exercise, on the other hand, serves as a potent modulator of cellular redox balance, which improves mitochondrial efficiency, reduces lipid peroxidation, and enhances antioxidant capacity [14–18].

Although each of these factors—aging, sex, and exercise—has been independently studied in relation to ferroptosis, little is known about their potential interactive effects, particularly in the context of skeletal muscle. Skeletal muscle is highly metabolically active tissue that is particularly sensitive to redox imbalances [19], iron accumulation [20], and oxidative stress [21]. Notably, iron accumulation activates multiple signaling pathways that induce ferroptosis, leading to a decline in muscle mass and fiber diameter and thereby impairing skeletal muscle function—a key pathological mechanism contributing to sarcopenia [6]. Those factors are profoundly influenced by aging, biological sex, and physical activity, so examining them is suitable for investigating ferroptosis dynamics. Furthermore, comprehensive, multi-dimensional analyses that concurrently assess ferroptosis at the histological, protein, and gene expression levels are lacking. In particular, the differential effects of exercise modalities—and their interactions with biological sex and aging—remain underexplored.

To address those knowledge gaps, we used a natural aging mouse model to investigate how aging, biological sex, and exercise modality interact to regulate ferroptosis in skeletal muscle. A multi-layered approach was used to assess histological features (e.g., fibrosis, iron accumulation), protein expression (e.g., GPX4, SLC7A11, p-AMPK), and gene-level markers (e.g., ACSL4, GCLC, POR). We placed special emphasis on delineating sex-specific responses to voluntary versus forced exercise in aged mice. This integrative framework was intended to provide novel insights into the regulatory mechanisms of ferroptosis during aging and inform the future development of sex- and modality-specific exercise interventions targeting ferroptosis-related muscle deterioration.

## Methods

### Animals and Experimental Design

The experimental animals were C57BL/6N mice that were randomly divided into 8 groups: young female control (N=5, 7 months old, FYC), young male control (N=5, 7 months old, MYC), old female control (N=6, 17 months old, FOC), old male control (N=6, 17 months old, MOC), old female voluntary exercise (N=6, 17 months old, FOW), old male voluntary exercise (N=6, 17 months old, MOW), old female forced exercise (N=6, 17 months old, FOT), and old male forced exercise (N=6, 17 months old, MOT). The FOW and MOW groups ran (5 times/week for 8 weeks) on a voluntary wheel (MAN86130, Lafayette Instrument Company, Lafayette, IN, USA). The FOT and MOT groups performed forced exercise (5 times/week for 8 weeks) on a treadmill (JD-A-07RA5, BS Technolab INC., Seoul, Korea) (Figure 1). During the first three weeks, the mice were acclimatized to their environment and underwent baseline testing (e.g., weekly voluntary wheel running distance). All mice were fed a standard chow diet (Teklad Global 2018, Envigo Inc.). Room temperature (22 ± 2 °C), humidity (50 ± 5%), and light (12/12 h light/dark cycle) were controlled, and the mice had *ad libitum* access to food, water, and activity. All the procedures followed in this experiment were approved by the Institutional Animal Care and Use Committee of Hanyang University (HYU 2021-0239A).

**Figure 1.**
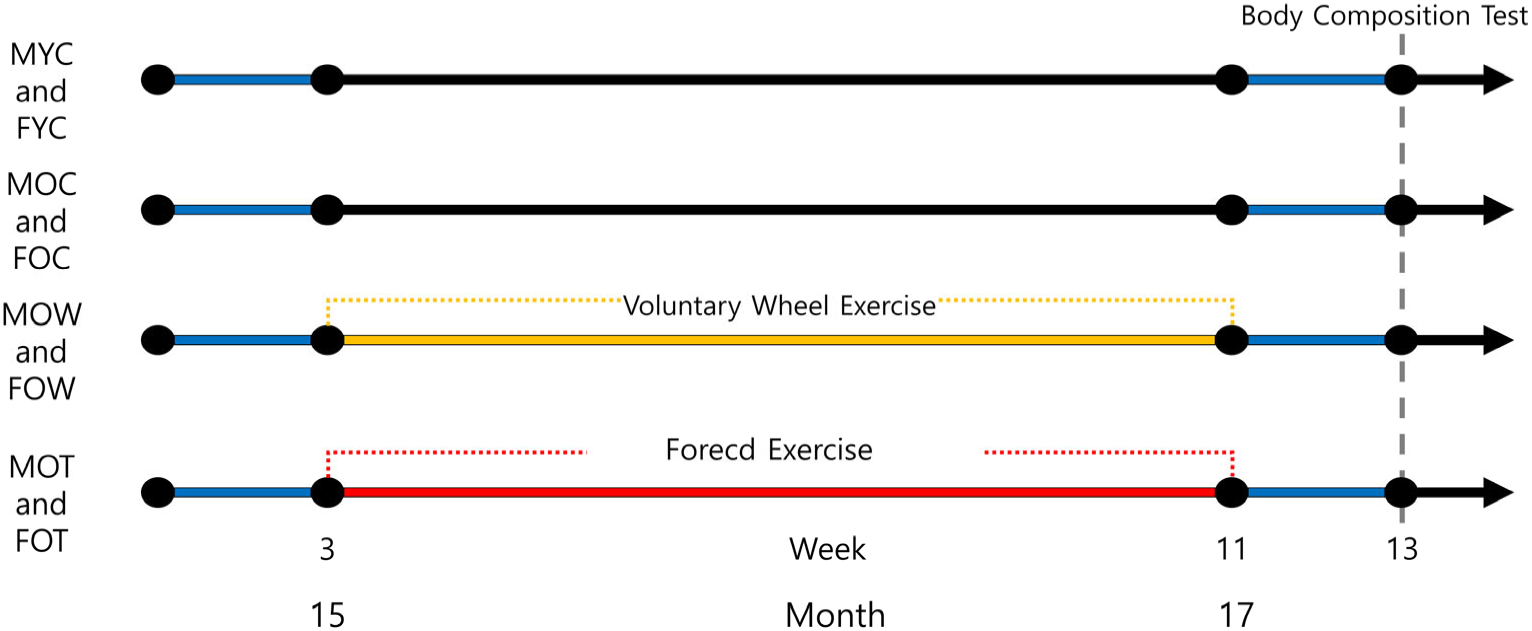
Schematic overview of the experimental study design.

The FOT and MOT mice were subjected to forced high-intensity interval training (HIIT), with running distances individualized based on the baseline testing. Each HIIT session consisted of five 5-minute intervals, totaling 25 minutes. During the first four weeks, the duration of low-intensity and high-intensity exercise was set at 2 minutes and 3 minutes, respectively. In the following four weeks, the duration changed to 1 minute of low-intensity and 4 minutes of high-intensity exercise. The low-intensity exercise started at 2 m/min, with an increase of 1 m/min every two weeks, and the high-intensity exercise started at 4 m/min, with an increase of 2 m/min each week (Table 1). After the 8-week intervention, the total running distance in old mice did not differ significantly between voluntary wheel running and HIIT exercise, suggesting similar overall running volume between modalities (Supplementary Document 1).

**Table 1.** HIIT Protocol.

| Week | Frequency | Time (min) |  | Intensity (m/min) |  | Set | Etc. |
| --- | --- | --- | --- | --- | --- | --- | --- |
|  |  | Low | High | Low | High |  |  |
| 1 |  | 2 | 3 | 2 | 4 |  |  |
| 2 |  | 2 | 3 | 2 | 6 |  |  |
| 3 |  | 2 | 3 | 3 | 8 |  | Run at 5 m/min speed with a |
| 4 | 5/week | 2 | 3 | 3 | 10 | 5 | 10-minute rest before and after |
| 5 |  | 1 | 4 | 4 | 12 |  | each exercise block for warm up |
| 6 |  | 1 | 4 | 4 | 14 |  | and clod down. |
| 7 |  | 1 | 4 | 5 | 16 |  |  |
| 8 |  | 1 | 4 | 5 | 18 |  |  |

### Physical Phenotype Test

All mice underwent tests before euthanasia. As previously described [22], the tests were walking speed, endurance, physical activity, grip strength, and body composition.

#### Rota-rod test

Walking speed was evaluated using the Rota-rod test. During the first three days, an adaptation period was implemented, consisting of a pre-test exercise at 5 rpm for 1 minute, once daily. The official test was performed in acceleration mode, with the speed increasing gradually from 5 to 50 rpm over 5 minutes (Model 76-0770, Harvard Apparatus Inc., Holliston, MA, USA). The time taken for the mouse to fall from the device (latency to fall) was recorded. Each mouse underwent three trials with a 10-minute rest interval between tests, and the best result among the three trials was used as the final outcome measure.

#### Treadmill test

Endurance capacity was assessed using a treadmill test. An adaptation phase, during which mice ran at a speed of 5 cm/s for 5 minutes on a 0-degree incline, was conducted once daily for three days prior to the official test. For the official test, the treadmill speed started at 5 cm/s and increased by 1 cm/s every 20 seconds, with the incline maintained at 0 degrees. The test concluded when the mouse touched the shock pad (set at 0.3 mA) three times.

#### Voluntary wheel test

Physical activity was evaluated using the voluntary wheel test. Running distance was measured with a voluntary wheel (MAN86130, Lafayette Instrument Company, Lafayette, IN, USA), where each rotation corresponded to a distance of 0.4 m. The average running distance over a 5-day period was recorded for each mouse.

#### Inverted-cling grip test

Grip strength was evaluated using the inverted-cling grip test. An adaptation phase was conducted over three days, with mice undergoing one adaptive trial per day. During the official test, each mouse was placed at the center of a wire mesh screen, and a stopwatch was started. The screen was gradually inverted over 2 seconds, positioning the mouse head-down, and held 40–50 cm above a padded surface. The time until the mouse fell was recorded. This procedure was repeated three times with 10-minute intervals between trials, and the highest recorded time was used as the final result.

#### Body composition

The mice were anesthetized with 40 mg/kg ketamine and 0.8 mg/kg medetomidine before undergoing a body composition analysis using an InAlyzer dual-energy X-ray absorptiometry (DEXA) system (Micro Photonics Inc., PA, USA). The measured parameters were the total mass (g), lean mass ratio (%), fat mass (g), lean mass (g), fat mass in tissue (%), and fat mass ratio (%) [23].

### Preservation and Preparation of Skeletal Muscle

The anesthetized (40 mg/kg ketamine and 0.8 mg/kg medetomidine) mice were euthanized by cervical dislocation [24]. Following euthanasia, the quadriceps (QD) muscles were carefully dissected. The left QD was fixed in 10% formalin (HT501128, Sigma,) for 24 hours, then dehydrated and embedded in paraffin for subsequent histological analyses: hematoxylin and eosin (H&E) staining, Prussian blue staining, Masson’s trichrome staining, and immunohistochemistry (IHC). The right QD was snap-frozen in liquid nitrogen and stored at –80°C for subsequent protein and gene expression analyses: biochemical assays, western blotting (WB), and quantitative reverse transcription-polymerase chain reaction (RT-qPCR).

### H&E staining

As previously described, rehydrated tissue sections (5 μm) were stained with ClearView™ hematoxylin (MA0101010, StatLab, McKinney, TX, USA) and ClearView™ eosin (MA0101015, StatLab, McKinney, TX, USA), and then the sections were dehydrated, cleared, and mounted with neutral resin [8, 24]. Images were captured using a slide scanner (Axio Scan.Z1, Zeiss) with a 20× objective lens. For each mouse, one QD section was randomly selected and photographed to obtain five images, each covering an area of 0.271 mm². The mean of the five images was calculated to represent the cross-sectional area (CSA) of the QD for that mouse. Fiber CSA was calculated automatically using Cellpose, an open-source deep learning–based segmentation tool [24].

### Prussian Blue Staining

Rehydrated tissue sections (5 μm) were stained using a Prussian blue staining kit (G1422, Beijing Solarbio Science & Technology Co., Ltd.) as previously described [8, 24]. Images were acquired on an Axio Scan. Z1 slide scanner (Zeiss) equipped with a 20× objective lens. For each mouse, one QD section was randomly selected and photographed to capture five images, each covering an area of 0.271 mm². The mean value of the five images is used to represent iron accumulation for that mouse. Images were analyzed automatically using DeepLIIF, an open-source deep learning–based segmentation [24].

### Masson’s Trichrome Staining

As previously described, rehydrated tissue sections (5 μm) were stained using a Masson’s trichrome staining kit (G1340, Beijing Solarbio Science & Technology Co., Ltd.) [8, 24]. A slide scanner (Axio Scan. Z1, Zeiss) fitted with a 20× objective was used to capture all tissue images. For each mouse, one QD section was randomly selected and photographed to obtain five images, each covering an area of 0.271 mm². The mean of the five images was calculated to represent the collagen fiber content in the QD for that mouse. Images were analyzed automatically using DeepLIIF [24].

### Immunohistochemistry

The IHC protocol was based on our prior study [8] with some modifications. Briefly, paraffin-embedded tissue sections (6 μm) were deparaffinized and rehydrated. Heat-induced antigen retrieval was conducted using 0.01 mol/L sodium citrate buffer (pH 6.0, C1010, Beijing Solarbio Science & Technology Co., Ltd.) in a microwave oven at 95–100°C for 20 minutes.

Following antigen retrieval, the sections were treated with 3% hydrogen peroxide (4104-4400, DAE JUNG) for 15 minutes twice to eliminate endogenous peroxidase activity. Afterward, they were blocked with blocking buffer (5% bovine serum albumin in phosphate-buffered saline) for 30 minutes at room temperature. The sections were then incubated with the primary antibody (Table 2) overnight (16–18 hours) at 4°C, followed by incubation with the secondary antibody (Table 2) for 60 minutes at room temperature. Finally, the sections were developed using a Pierce™ DAB substrate kit (#34002, Thermo Scientific).

**Table 2.** Primary and Secondary Antibodies.

| Antibodies | Source | Vender | Catalog | Dilution (WB) | Dilution (IHC) |
| --- | --- | --- | --- | --- | --- |
| P53 | Rabbit | Cell signaling | #30313 | 1:500 | - |
| SLC7A11 | Rabbit | Abcam | ab175186 | 1:1000 | - |
| GAPDH | Rabbit | Cell signaling | #5174 | 1:6000 | - |
| AMPK | Rabbit | Cell signaling | #5831 | 1:500 | - |
| p-AMPK<sup>Thr172</sup> | Rabbit | Cell signaling | #2535 | 1:400 | - |
| GPX4 | Rabbit | Abcam | ab125066 | 1:1000 | - |
| Tfr1 | Rabbit | Abcam | ab84036 | 1:500 | - |
| NCOA4 | Rabbit | Invitrogen | #PA5-115626 | 1:500 | - |
| Rab7 | Rabbit | Cell signaling | #9376 | 1:500 | - |
| 4-HNE | Rabbit | Abcam | ab46545 | 1:1000 | 1:200 |
| MDA | Mouse | Abcam | ab243066 | 1:250 | 1:50 |
| Goat anti Rabbit | Goat | Invitrogen | #G-21234 | 1:5000 | 1:500 |
| m-IgG Fc | - | SANTA | sc-525409 | 1:5000 | 1:25 |

All images were obtained using an Axio Scan. Z1 (Zeiss) slide scanner operating with a 20× objective lens. For each mouse, one QD section was randomly selected and photographed to obtain five images, each covering an area of 0.271 mm². The mean of the five images was used to quantify antibody expression in the QD for that mouse. Images were analyzed automatically using DeepLIIF, an open-source deep learning–based segmentation tool [24].

### WB Immunoblotting

Following the WB protocol of a prior study [23] with some modifications, each QD was lysed using an EzRIPA lysis kit (WSE-7420, ATTO, Japan). The protein concentration was measured using a Pierce™ bicinchoninic acid protein assay kit (#23225, Thermo Scientific, Waltham, MA, USA). Equal amounts of protein (35 µg) were separated via sodium dodecyl sulfate–polyacrylamide gel electrophoresis and transferred onto nitrocellulose membranes (#1620112, Bio-Rad Laboratories, Hercules, CA, USA).

The membranes were blocked with 5% non-fat milk dissolved in Tris-buffered saline with Tween-20 (10 mM Tris, 150 mM NaCl, and 0.1% Tween-20; pH 7.6) for 1.5 hours at room temperature, followed by overnight incubation (16–18 hours) at 4°C with the primary antibodies (Table 2). Afterward, the membranes were incubated with horseradish peroxidase–conjugated secondary antibodies (Table 2) for 1.5 hours at room temperature. Protein expression levels were normalized to GAPDH, which served as an internal control. The targeted protein bands were quantified by densitometry using ImageJ software (National Institutes of Health, USA).

### Iron Assay

Total iron content in skeletal muscle was measured using a colorimetric iron assay kit according to the manufacturer’s protocol (Beijing Solarbio Science & Technology Co., Ltd., #BC4355). In brief, muscle tissues were homogenized in the provided extraction buffer and centrifuged at 12,000 ×g for 10 min at 4°C. The supernatant was collected, and iron was reduced to the ferrous (Fe²⁺) form with the supplied reducer. Subsequently, the ferrous iron reacted with the chromogen reagent to form a stable-colored complex. After incubation at room temperature for 30 min, absorbance was measured at 593 nm using a microplate reader. Iron concentration was determined from a standard curve and normalized to the protein content (µmol/µg protein).

### Malondialdehyde (MDA) Assay

MDA levels were determined using the thiobarbituric acid (TBA) method with a commercially available assay kit (Beijing Solarbio Science & Technology Co., Ltd., #BC0025). Skeletal muscle tissues were homogenized in cold phosphate-buffered saline (PBS) and centrifuged at 10,000 ×g for 10 min at 4°C. The supernatant was then mixed with TBA reagent and incubated in a 95°C water bath for 40 min to form the MDA-TBA adduct. After cooling to room temperature and centrifugation to remove precipitates, absorbance was measured at 532 nm using a microplate reader. MDA concentration was calculated from a standard curve and expressed as nmol/g protein.

### Oxidized Glutathione (GSSG) Assay

GSSG levels were measured using a glutathione assay kit based on the enzymatic recycling method (Beijing Solarbio Science & Technology Co., Ltd., #BC1185). Tissues were homogenized in cold metaphosphoric acid or the kit-provided extraction buffer and centrifuged at 10,000 ×g for 10 min at 4°C. The resulting supernatant was treated with a GSH-scavenging reagent to remove reduced glutathione (GSH), allowing selective measurement of GSSG. The remaining GSSG was then reduced to GSH by glutathione reductase in the presence of NADPH, and the newly formed GSH reacted with 5,5’-dithiobis-(2-nitrobenzoic acid) (DTNB) to generate the yellow-colored product (TNB). Absorbance was recorded at 412 nm using a microplate reader, and GSSG levels were calculated from a standard curve and expressed as µg/ml.

### Reduced Glutathione (GSH) Assay

GSH levels were measured using a glutathione assay kit based on the enzymatic recycling method, according to the manufacturer’s protocol (Beijing Solarbio Science & Technology Co., Ltd., #BC1175). Skeletal muscle tissues were homogenized in cold metaphosphoric acid or the kit-provided extraction buffer and centrifuged at 10,000 ×g for 10 min at 4°C. The supernatant was collected and directly used for GSH quantification. GSH reacted with DTNB to produce TNB, and signal amplification was achieved using glutathione reductase and NADPH. Absorbance was measured at 412 nm, and GSH concentration was determined from a standard curve and expressed as µg/ml.

### RT-qPCR

Each QD was homogenized in TRIzol (#15596018, Thermo Scientific) and briefly centrifuged at maximum speed, and then the supernatant was transferred to a new tube. Chloroform (#288306, Sigma) was added to the tube, which was then securely capped and thoroughly mixed by shaking, followed by a 5-minute incubation. Each sample was centrifuged at 12,000 × g for 15 minutes at 4°C, resulting in separation into three layers: a lower phenol-chloroform phase, an interphase, and a colorless upper aqueous phase. The RNA-containing aqueous phase was carefully transferred to a new tube by angling the tube to 45° and gently pipetting out the solution.

Isopropanol (#563935, Sigma) was added to the aqueous phase, mixed thoroughly, and incubated for 10 minutes. Each sample was then centrifuged at 12,000 × g for 10 minutes at 4°C, which produced a white, gel-like RNA pellet at the bottom of the tube. The supernatant was carefully discarded, and the pellet was washed by resuspending it in 75% ethanol, vortexing it briefly, and centrifuging it at 7,500 × g for 5 minutes at 4°C. This washing process was repeated 2–3 times. The RNA pellet was then air-or vacuum-dried for 5–10 minutes and resuspended in DEPC-treated water (#R0601, Thermo Scientific). After its purity and concentration were measured with a NanoDrop™ One/OneC microvolume UV-Vis spectrophotometer (Thermo Fisher Scientific, Cleveland, OH, USA), it was stored at –80°C.

Quantitative PCR was performed using a high-capacity RNA-to-cDNA™ kit (#4387406, Thermo Scientific) with PowerUp™ SYBR™ Green PCR master mix (#A25742, Thermo Scientific). Each reaction was set up in a total volume of 20 µL, containing 4 µL of diluted cDNA (equivalent to 50 ng of starting RNA), 10 µL of 2X SYBR Green PCR master mix, 0.5 µM of each forward and reverse primer, and 5 µL of nuclease-free water. The primer sequences used for amplification are listed in Table 3.

**Table 3.**
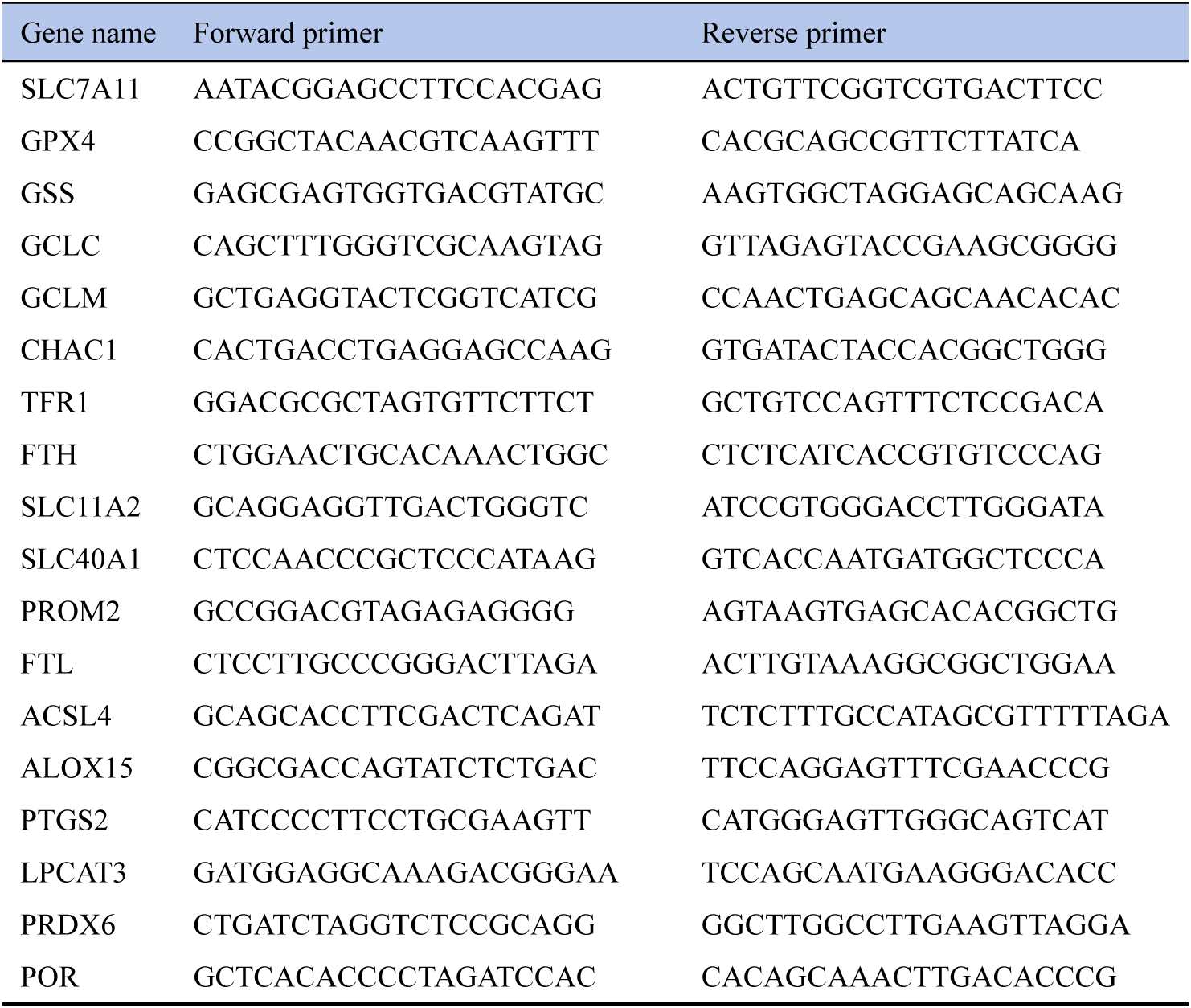
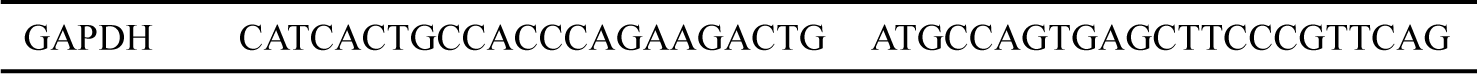
Primer Sequences.

All reactions were performed in triplicate, and the average threshold cycle (Ct) values were calculated. Relative gene expression was quantified using the 2^^−ΔΔCt^ method [25], with GAPDH serving as the endogenous control. The ΔCt was calculated as the difference between the Ct value of the target gene and the Ct value of the housekeeping gene, and ΔΔCt was determined by comparing the ΔCt of the experimental sample with that of the control sample.

### Statistical Analysis

Statistical analyses were conducted using GraphPad Prism software (version 9.4). To assess the effects of aging (young vs. old), sex (male vs. female), and exercise type (voluntary vs. forced), a two-way ANOVA with Bonferroni’s post-hoc test was applied. Outliers were checked using the ROUT method with Q = 1% in GraphPad Prism, and no outliers were identified or excluded. The linear relationship between protein expression levels and physical phenotypes was evaluated using Pearson correlation analysis. An R² value greater than 0.4 was considered indicative of a meaningful correlation, while an R² value exceeding 0.5 was regarded as evidence of a strong correlation [26, 27]. All results are expressed as the mean ± standard error of the mean (SEM). Statistical significance was set at p < 0.05, with asterisks indicating the following levels of significance: *p < 0.05, **p < 0.01, ***p < 0.001, and ****p < 0.0001.

## Results

### 3.1 Aging Induces Sex-Specific Alterations in Physical Phenotype

Aging led to a significant increase in body weight, primarily attributable to elevated fat mass and percentage (fat mass in tissue) in both male and female mice (Figure 2A and F). Although weight gain occurred in both sexes, aged females (FOC) exhibited lower body weight than aged males (MOC), likely reflecting sex-based differences in fat accumulation (Figure 2C and F). Aging also significantly reduced grip strength, endurance, and physical activity. Notably, endurance performance displayed a sex-specific response, suggesting differential physiological adaptations to aging between males and females (Figure 2G–J).

**Figure 2.**
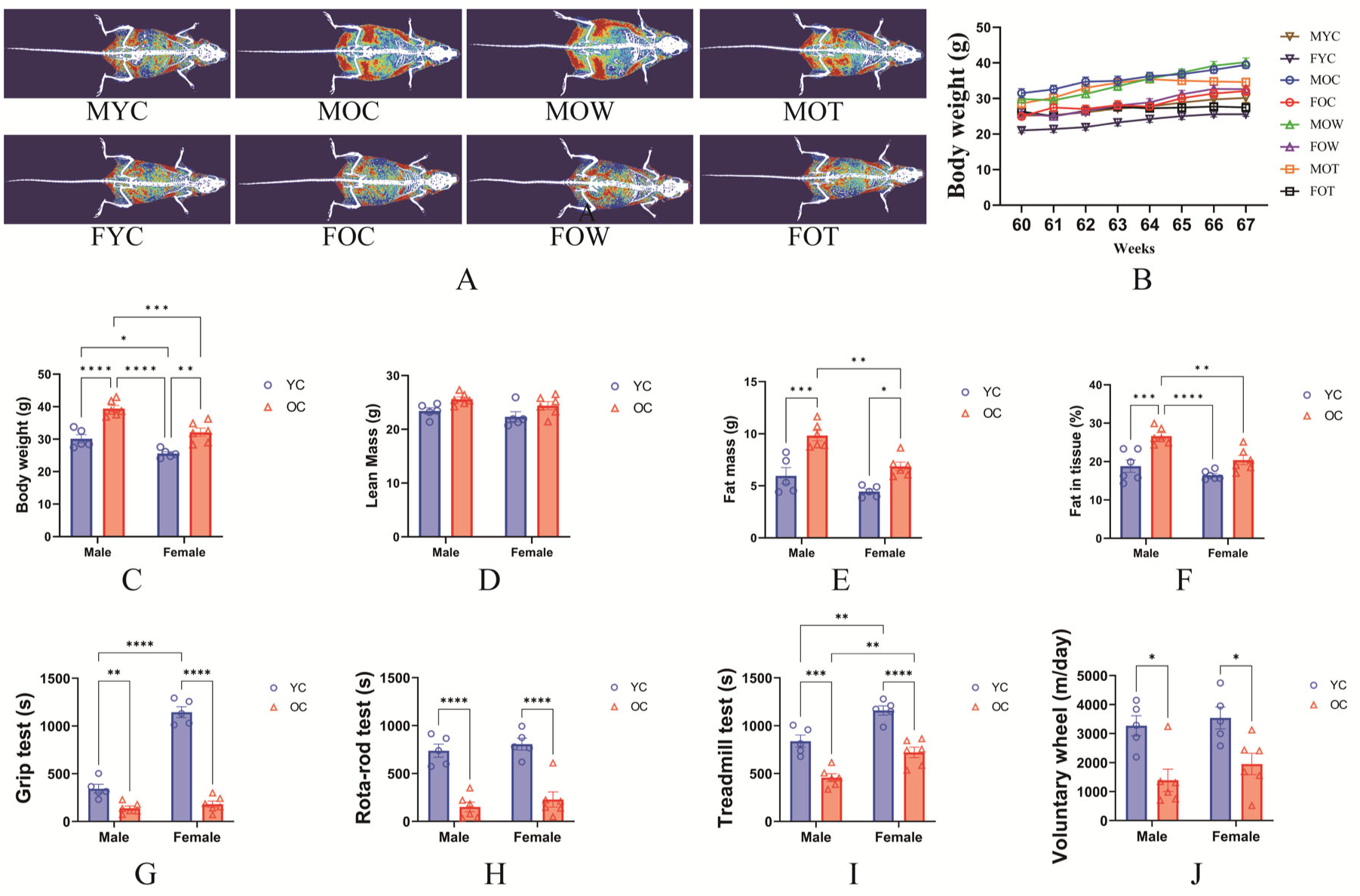
(A) Representative DEXA scan images, with skeletal muscle, fat tissue, and bone shown in blue, red, and white, respectively. (B) Weekly body weight changes (group × weeks) (60–67 weeks). (C–F) Quantitative data on final body weight (g), lean mass (g), fat mass (g), and fat mass in tissue (%), respectively, were acquired by DEXA scanning. (H–J) Quantitative data from the grip strength test, Rota-rod test (walking speed test), treadmill test (endurance capacity test), and voluntary wheel test (physical activity test). Significant differences are denoted by an asterisk: P<0.05 (*), P<0.01 (**), P<0.001 (***), and P<0.0001 (****). All values are presented as the mean ± SEM.

### 3.2 Regulation of Ferroptosis-related Histological Markers by Aging Is Minimally Affected by Sex

Both aging and elevated body weight have been implicated in promoting ferroptosis across various tissues and organ systems [1, 8]. However, the regulatory role of the interaction between aging and sex remains unclear. Therefore, we conducted a histological analysis of QD muscle tissue.

Canonical histological indicators of ferroptosis include inflammatory cell infiltration [28, 29], fibrosis [30], ferric iron accumulation [31], and increased oxidative stress [29, 30]. Our data reveal that aging did not induce inflammatory cell infiltration in skeletal muscle (Figure 3A and B). In contrast, aging—regardless of sex—significantly increased skeletal muscle fibrosis and ferric iron deposition and elevated the expression of oxidative stress markers, particularly 4-hydroxynonenal (4-HNE) and malondialdehyde (MDA) (Figure 3D, E, G, H, G, K, M, and N). Collectively, these findings suggest that histological markers associated with ferroptosis are primarily regulated by aging, with no differences observed between sexes.

**Figure 3.**
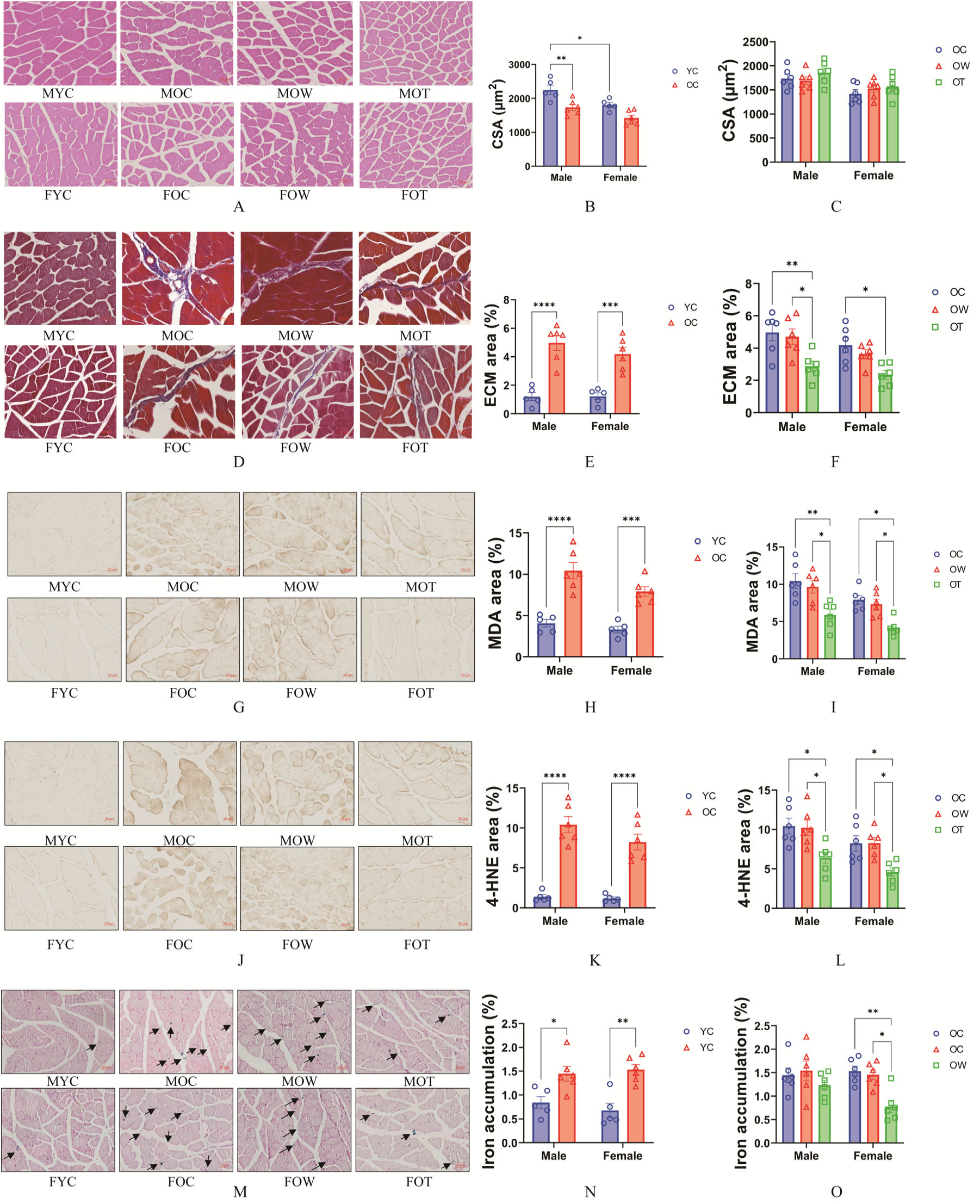
(A) Representative images showing H&E staining of QD. (B–C) Cross sectional areas of QD. (D) Representative images showing Masson’s trichrome staining of QD. (E–F) Skeletal muscle collagen area in QD. (G) Representative images showing MDA expression in QD. (H–I) The expression of MDA in QD. (J) Representative images showing 4-HNE expression in QD. (K–L) The expression of 4-HNE in QD. (M) Representative images showing Prussian blue staining of QD, with black rows indicating regions of ferric iron (Fe³⁺) accumulation. (N–O) Ferric iron accumulation in QD. Significant differences are denoted by an asterisk: P<0.05 (*), P<0.01 (**), P<0.001 (***), and P<0.0001 (****). All values are presented as the mean ± SEM.

### 3.3 Aging and Sex Interact to Regulate Ferroptosis-Related Protein Markers

Although the histological indicators of ferroptosis appear to be primarily influenced by aging, independent of sex, we next examined whether this pattern extends to key ferroptosis-related protein markers. WB analyses revealed that aging significantly decreased the expression of GPX4, SLC7A11, and phosphorylated AMPK^Thr172^, with no observable sex differences (Figure 4A, C, and F). Conversely, aging markedly increased the expression of the oxidative stress makers MDA and 4-HNE, again without sex-specific differences (Figure 4J and K).

**Figure 4.**
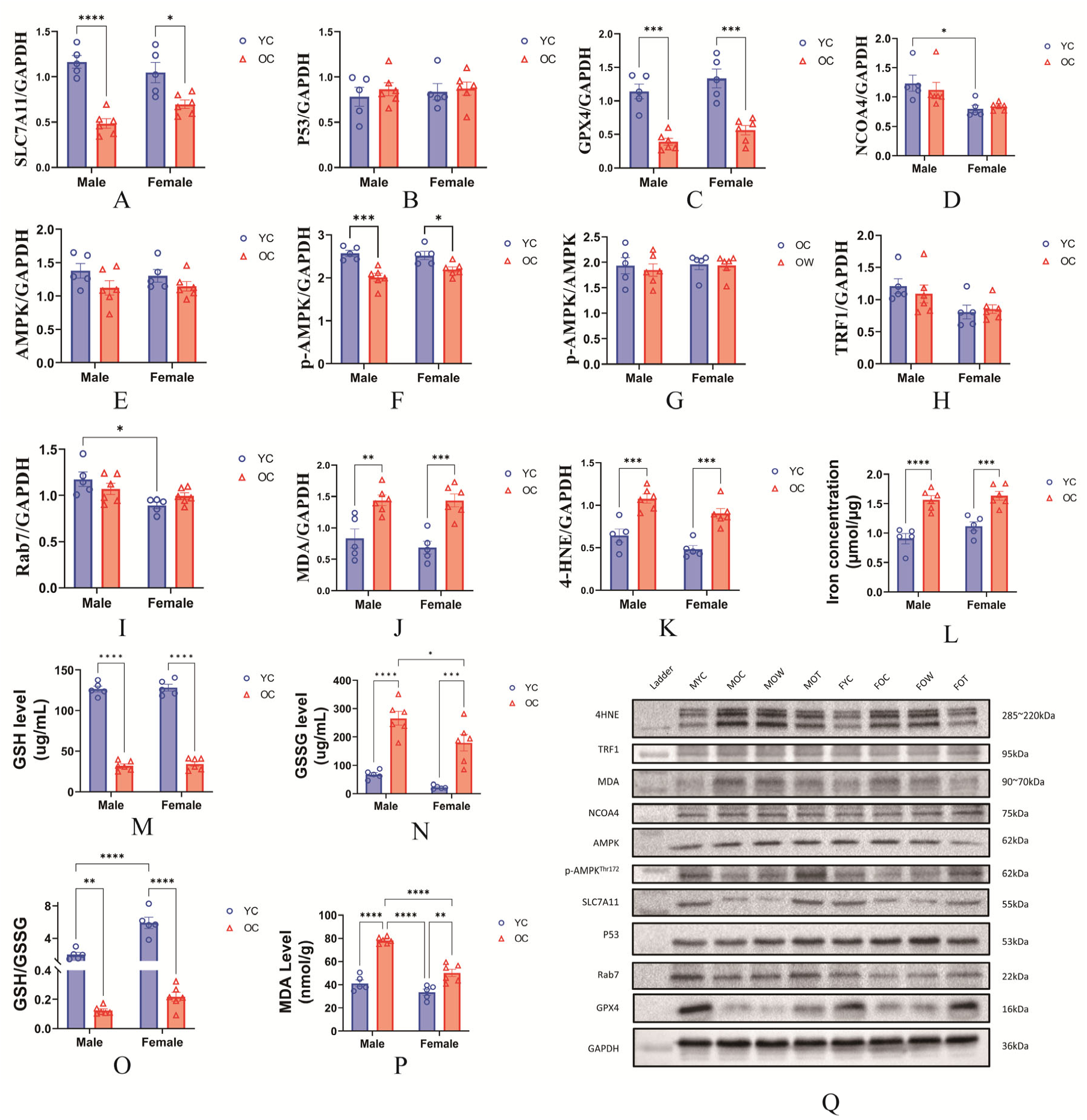
(A) Solute carrier family 7 member 11 (SLC7A11). (B) P53. (C) Glutathione peroxidase 4 (GPX4). (D) Nuclear receptor coactivator 4 (NCOA4). (E) AMP-activated protein kinase (AMPK). (F) Phospho-AMPKα (Thr172). (G) p-AMPK^Thr172^/AMPK ratio. (H) Transferrin receptor 1 (Tfr1). (I) Ras-related protein Rab-7a (Rab7). (J) Malonaldehyde (MDA). (K) 4-Hydroxynonenal (4-HNE). (L) Fe^3+^ concentration in skeletal muscle. (M) Glutathione (GSH) concentration in skeletal muscle. (N) Glutathione disulfide (GSSG) concentration in skeletal muscle. (O) GSH/GSSG ratio. (P) MDA concentration in skeletal muscle. (Q) Representative WB image. Significant differences are denoted by an asterisk: P<0.05 (*), P<0.01 (**), P<0.001 (***), and P<0.0001 (****). All values are presented as the mean ± SEM.

To further validate these observations, we conducted enzymatic assay tests. Although the GSH levels and GSH/GSSG ratio did not display sex specificity, GSSG expression exhibited a significant interaction effect, with aged males displaying higher levels than aged females (Figure 4M, N, and O). Additionally, MDA concentrations were affected by the interaction between aging and sex, with the MOC group exhibiting higher levels than the FOC group (Figure 4P). Lower levels of GSSG and MDA are indicative of reduced oxidative stress, suggesting a decreased vulnerability to ferroptosis [32, 33].

Moreover, the Fe³⁺ concentration in skeletal muscle increased exclusively with aging and was not affected by sex (Figure 4L), further supporting the histological findings. Collectively, these results indicate that aging independently modulates the expression of most ferroptosis-related protein markers, with GSSG and MDA concentrations being jointly regulated by the interaction of aging and sex.

### 3.4 Aging Regulates Ferroptosis-related Gene Expression in a Sex-specific Manner

Our WB analyses indicate that the expression of certain ferroptosis protein markers is modulated by both aging and sex. To further elucidate the transcriptional mechanisms underlying those observations, we used RT-qPCR to examine the mRNA expression profiles of key ferroptosis-associated genes (Figure 5).

**Figure 5.**
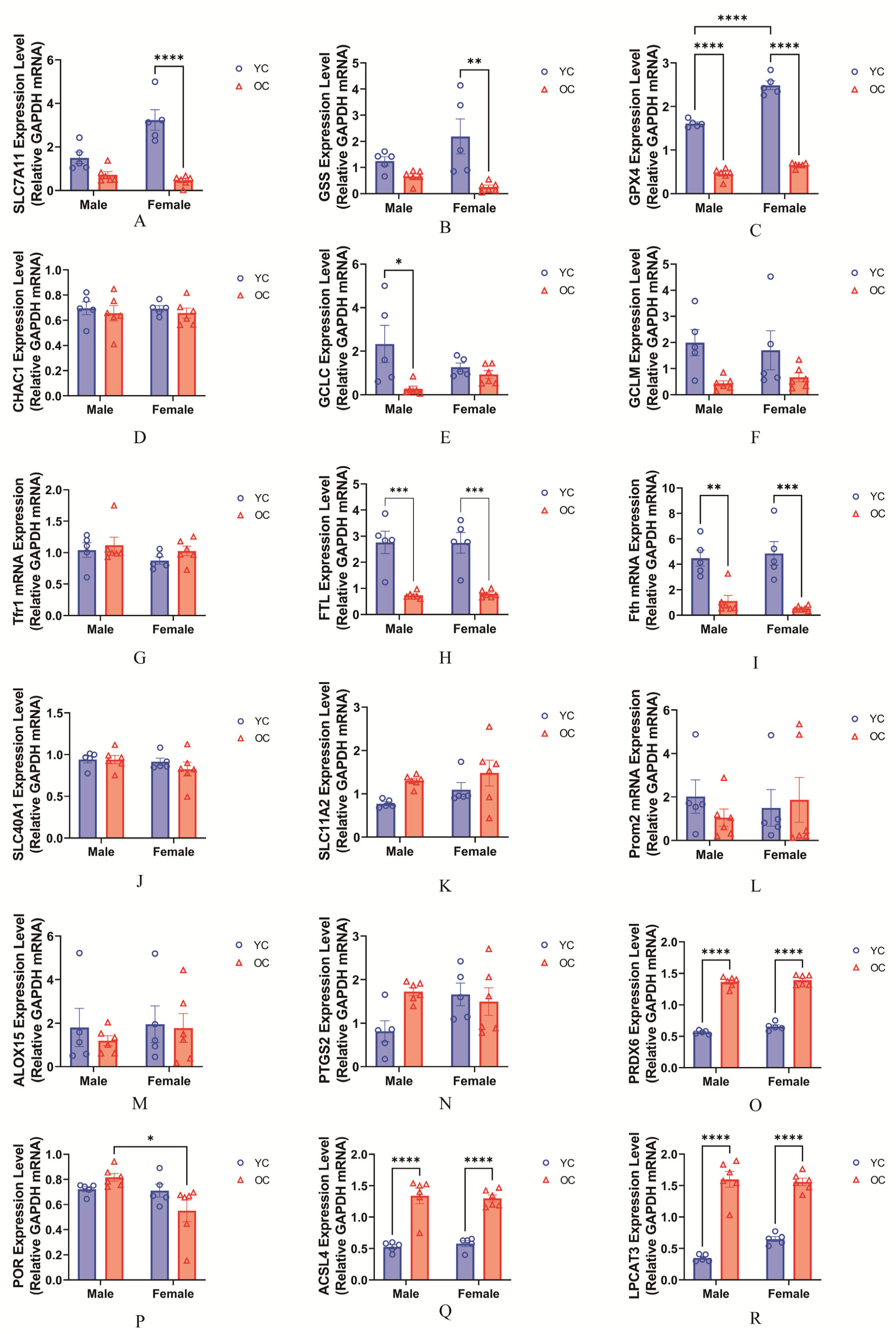
(A) Solute carrier family 7 member 11 (SLC7A11). (B) Glutathione synthetase (GSS). (C) Glutathione peroxidase 4 (GPX4). (D) CHAC glutathione-specific gamma-glutamylcyclotransferase 1 (CHAC1). (E) Glutamate-cysteine ligase catalytic subunit (GCLC). (F) Glutamate-cysteine ligase modifier subunit (GCLM). (G) Transferrin receptor 1 (Tfr1). (H) Ferritin light chain (FTL). (I) Ferritin heavy chain (FTH). (J) Solute carrier family 40 member 1 (SLC40A1). (K) Cationic amino acid transporter 2 (SLC7A2). (L) Prominin 2 (PROM2). (M) Arachidonate 15-lipoxygenase (ALOX15). (N) Prostaglandin-endoperoxide synthase 2 (PTGS2). (O) Peroxiredoxin 6 (PRDX6). (P) Cytochrome P450 oxidoreductase (POR). (Q) Acyl-CoA synthetase long chain family member (ACSL4). (R) Lysophosphatidylcholine acyltransferase 3 (LPCAT3). Significant differences are denoted by an asterisk: P<0.05 (*), P<0.01 (**), P<0.001 (***), and P<0.0001 (****). All values are presented as the mean ± SEM.

Aging exerted distinct sex-specific effects on gene expression, particularly in the amino acid (antioxidant) (Figure 5A–F) and lipid metabolism pathways (Figure 5M–R). In the amino acid pathway, aging significantly downregulated SLC7A11, GSS, GPX4, and GCLC gene expression (Figure 5A, B, C, and E). However, SLC7A11 and GSS were selectively reduced in aged females, whereas GCLC expression was selectively suppressed in aged males, indicating sexually dimorphic transcriptional regulation.

In contrast, the iron metabolism pathway showed minimal alterations, with only ferritin heavy chain expression significantly affected by aging (Figure 5I). In the lipid metabolism pathway, PRDX6, ACSL4, and LPCAT3 expression levels were consistently downregulated with aging in both sexes (Figure 5O, Q, and R). Notably, although POR expression was not significantly regulated by aging per se, its levels were markedly lower in aged females than in aged males, suggesting sex-specific regulatory control (Figure 5P). Taken together, these findings demonstrate that aging induces sex-dependent alterations in the transcriptional regulation of ferroptosis-related genes, with differential responses observed across metabolic pathways.

### 3.5 Interaction Between Exercise Modality and Sex Affects the Physical Phenotype of Aged Mice

In the aged state, the interaction between exercise modality and biological sex produced distinct effects on physical phenotype. Regarding body composition, forced exercise resulted in a more substantial reduction in body weight than voluntary exercise; however, the underlying mechanisms of this reduction differed by sex. In the FOT group, forced exercise significantly decreased both fat mass and tissue fat percentage, whereas in the MOT group, only fat mass was reduced, with fat content in tissue remaining unchanged (Figure 6A–D).

**Figure 6.**
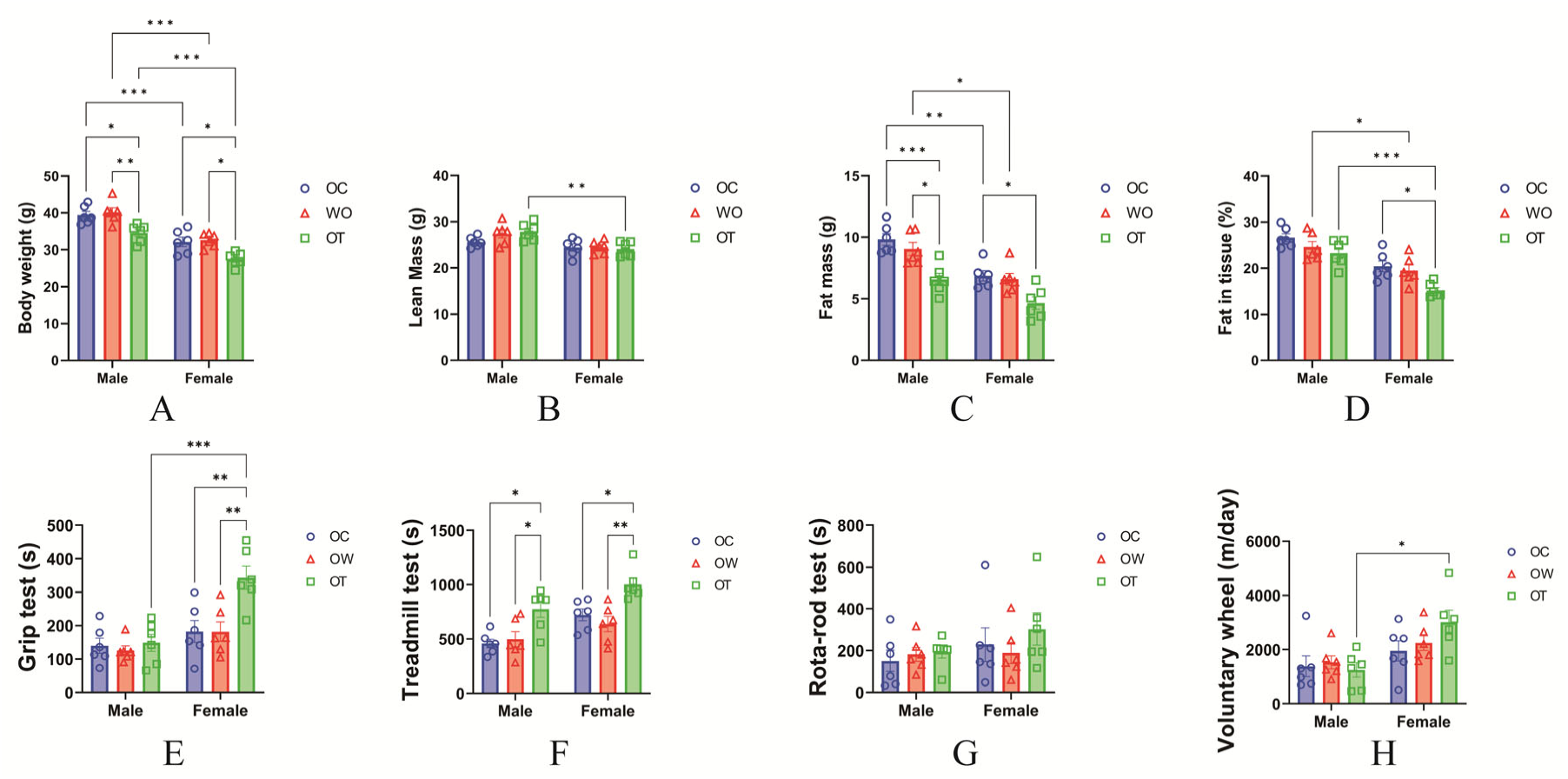
(A–D) Quantitative data on final body weight (g), lean mass (g), fat mass (g), and fat mass in tissue (%), respectively, were acquired by DEXA scanning. (E–H) Quantitative data from the grip strength test, Rota-rod test (walking speed test), treadmill test (endurance capacity test), and voluntary wheel test (physical activity test). Significant differences are denoted by an asterisk: P<0.05 (*), P<0.01 (**), P<0.001 (***), and P<0.0001 (****). All values are presented as the mean ± SEM.

Skeletal muscle function also exhibited sex-specific responses to exercise modality. Notably, forced exercise led to significant improvements in grip strength and voluntary physical activity in females, whereas those benefits were less pronounced in males (Figure 6E and F).

### 3.6 Interaction Between Exercise Modality and Sex Influences Histological Ferroptosis Markers in Aged Mice

To further investigate how the interaction between exercise modality and sex regulates ferroptosis markers in aged skeletal muscle, we conducted histological analyses of key ferroptosis-associated markers (Figure 3). Those results show that forced exercise significantly attenuated skeletal muscle fibrosis, as well as MDA and 4-HNE expression, without notable sex differences (Figure 3F, I, and L). Interestingly, Fe³⁺ accumulation exhibited a significant interaction between exercise type and sex. Specifically, a marked reduction in ferric iron levels was observed only in the forced exercise groups (Figure 3O), suggesting sex-dependent regulation under certain exercise conditions. In summary, with the exception of Fe³⁺ accumulation, no significant interaction between exercise modality and sex was observed in the histological indicators of ferroptosis.

### 3.7 In Aged Mice, the Expression of Ferroptosis-related Protein Markers Is Modulated by an Interaction Between Exercise Modality and Sex

Our histological analyses indicated a potential interaction between exercise modality and sex in the regulation of ferroptosis markers. To further elucidate that relationship, we conducted WB and biochemical assays to examine the regulatory effects of exercise modality and sex on ferroptosis-associated protein markers (Figure 7).

**Figure 7.**
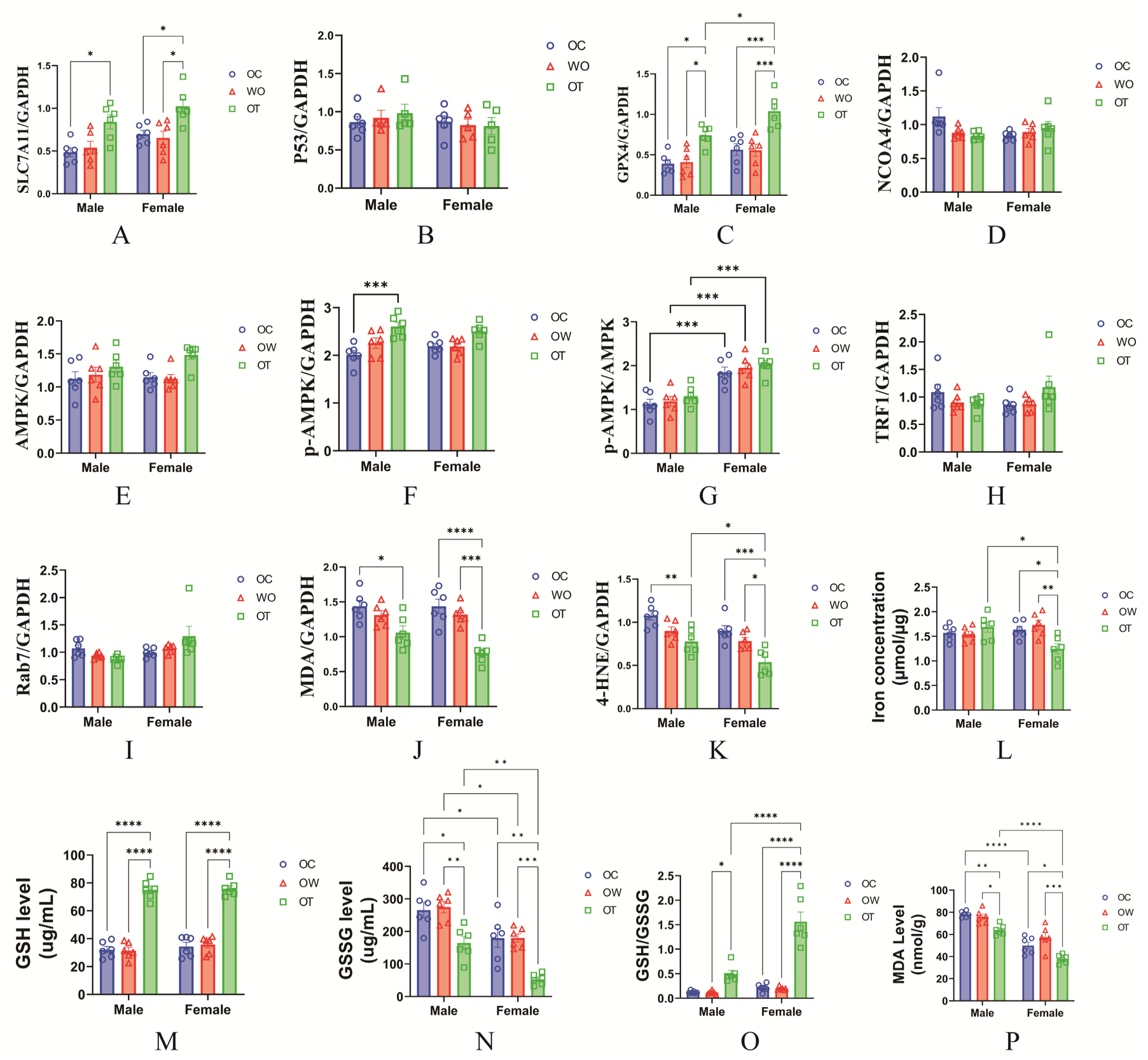
(A) Solute carrier family 7 member 11 (SLC7A11). (B) P53. (C) Glutathione peroxidase 4 (GPX4). (D) Nuclear receptor coactivator 4 (NCOA4). (E) AMP-activated protein kinase (AMPK). (F) Phospho-AMPKα (Thr172). (G) p-AMPK^Thr172^/AMPK ratio. (H) Transferrin receptor 1 (Tfr1). (I) Ras-related protein Rab-7a (Rab7). (J) Malonaldehyde (MDA). (K) 4-Hydroxynonenal (4-HNE). (L) Fe^3+^ concentration in skeletal muscle. (M) Glutathione (GSH) concentration in skeletal muscle. (N) Glutathione disulfide (GSSG) concentration in skeletal muscle. (O) GSH/GSSG ratio. (P) MDA concentration in skeletal muscle. Significant differences are denoted by an asterisk: P<0.05 (*), P<0.01 (**), P<0.001 (***), and P<0.0001 (****). All values are presented as the mean ± SEM.

The WB analyses revealed that only forced exercise significantly modulated the expression of ferroptosis protein markers. Specifically, the SLC7A11 (Figure 7A), p-AMPK (Figure 7F), and MDA (Figure 7J) proteins showed altered expression solely following forced exercise, independent of sex differences. Importantly, GPX4 (Figure 7C) and 4-HNE (Figure 7K) exhibited a clear interaction between exercise type and sex. Forced exercise increased GPX4 expression in both the MOT and FOT groups, with significantly higher levels observed in aged females than aged males. Conversely, 4-HNE expression was reduced in both groups by forced exercise, with aged female subjects exhibiting significantly lower levels than aged males.

Complementary biochemical assays supported those findings. The levels of Fe³⁺, GSSG, and MDA, as well as the GSH/GSSG ratio, all demonstrated significant interactions between exercise modality and sex. Notably, aged female mice in the forced exercise group exhibited significant reductions in Fe³⁺, GSSG, and MDA levels, along with an elevated GSH/GSSG ratio, all of which differed significantly from their male counterparts in identical conditions. Overall, these results indicate that, compared with aged male mice, aged female mice respond more robustly to forced exercise at the molecular level, particularly in the regulation of ferroptosis-related protein markers, suggesting sex-dependent susceptibility and adaptability to exercise-induced redox modulation in aging skeletal muscle.

### 3.8 Interaction Between Exercise Modality and Sex Regulates Ferroptosis-related Gene Expression in Aged Mice

Following confirmation from the WB analyses that ferroptosis-related protein markers are modulated by both exercise modality and sex in aged mice, we investigated transcriptional changes by using RT-qPCR to examine the expression of key genes associated with ferroptosis (Figure 8).

**Figure 8.**
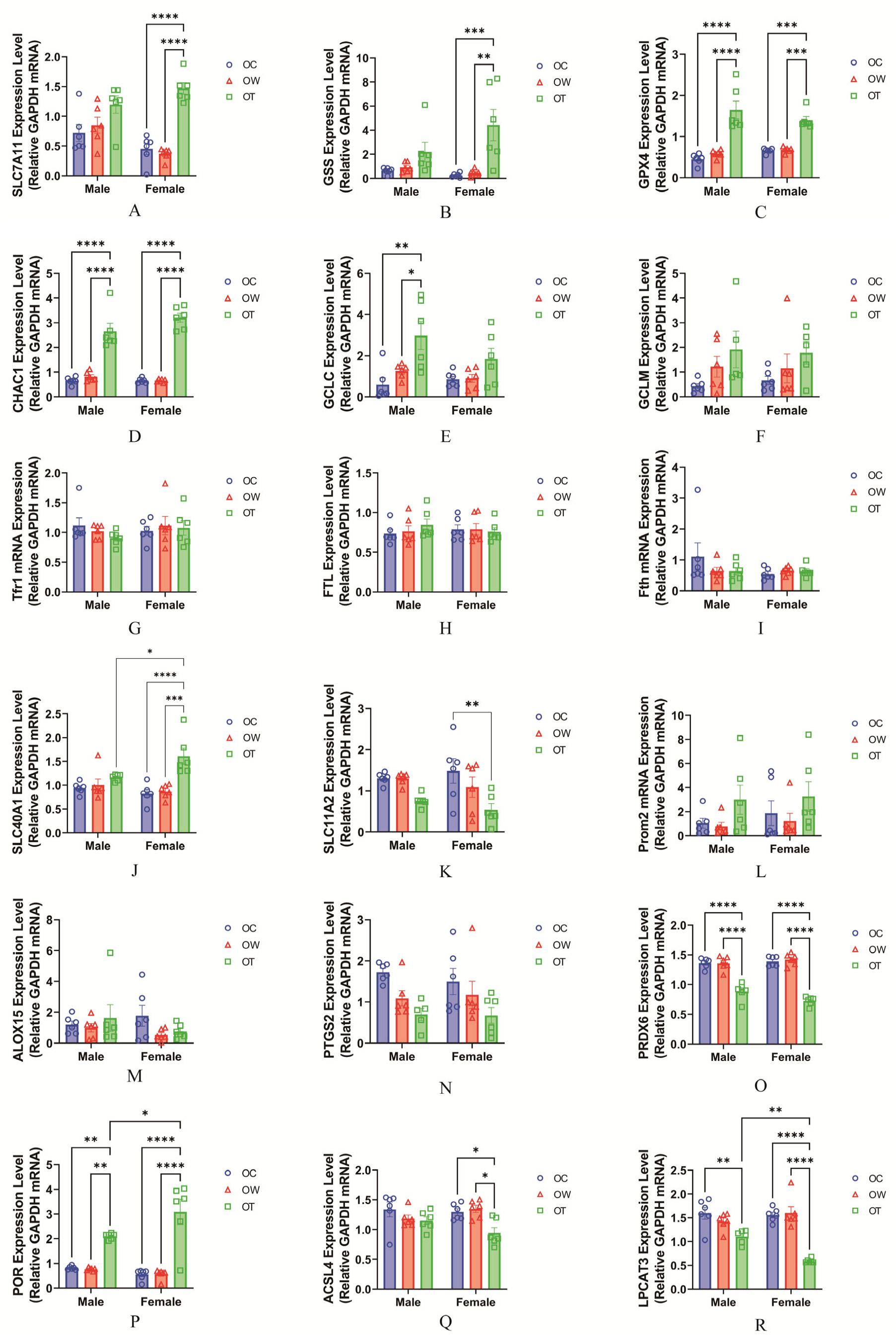
(A) Solute carrier family 7 member 11 (SLC7A11). (B) Glutathione synthetase (GSS). (C) Glutathione peroxidase 4 (GPX4). (D) CHAC glutathione-specific gamma-glutamylcyclotransferase 1 (CHAC1). (E) Glutamate-cysteine ligase catalytic subunit (GCLC). (F) Glutamate-cysteine ligase modifier subunit (GCLM). (G) Transferrin receptor 1 (Tfr1). (H) Ferritin light chain (FTL). (I) Ferritin heavy chain (FTH). (J) Solute carrier family 40 member 1 (SLC40A1). (K) Cationic amino acid transporter 2 (SLC7A2). (L) Prominin 2 (PROM2). (M) Arachidonate 15-lipoxygenase (ALOX15). (N) Prostaglandin-endoperoxide synthase 2 (PTGS2). (O) Peroxiredoxin 6 (PRDX6). (P) Cytochrome P450 oxidoreductase (POR). (Q) Acyl-CoA synthetase long chain family member (ACSL4). (R) Lysophosphatidylcholine acyltransferase 3 (LPCAT3). Significant differences are denoted by an asterisk: P<0.05 (*), P<0.01 (**), P<0.001 (***), and P<0.0001 (****). All values are presented as the mean ± SEM.

Our results reveal that several genes involved in the amino acid (antioxidant), iron metabolism, and lipid metabolism pathways are differentially regulated by exercise type and sex. In the amino acid pathway, SLC7A11, GSS, GPX4, CHAC1, and GCLC (Figure 8A–E) were significantly up- or downregulated following forced exercise. Specifically, GPX4 and CHAC1 were markedly altered in aged mice following forced exercise, irrespective of sex. In contrast, SLC7A11 and GSS were significantly modulated only in aged females, and GCLC expression was selectively altered in aged males. Within the iron metabolism pathway, only SLC40A1 and SLC11A2 responded significantly in aged females subjected to forced exercise (Figure 8J and K), with SLC40A1 levels notably elevated compared with their male counterparts. Regarding lipid metabolism, PRDX6, POR, ACSL4, and LPCAT3 (Figure 8O–R) displayed pronounced regulation after forced exercise. Forced exercise led to significant downregulation of PRDX6 in both sexes. Although POR and LPCAT3 were respectively up- and downregulated in a sex-independent manner after forced exercise, the magnitude of change was greater in females. Notably, ACSL4 was significantly downregulated only in aged females in the forced exercise group. Collectively, these data highlight a more robust transcriptional response to ferroptosis-related genes in aged female mice subjected to forced exercise than in their male counterparts.

### 3.9 Sex-specific Correlations Between Ferroptosis Signaling and Physical Phenotypes

To further investigate the relationship between physical phenotypes and ferroptosis, we performed correlation analyses between physical performance indicators and the protein expression levels of GPX4, MDA, and 4-HNE, which represent downstream regulatory nodes of the ferroptosis pathway and are widely recognized as critical modulators of ferroptosis susceptibility [2, 13, 34, 35].

In male mice, GPX4 expression was strongly positively correlated with walking speed (R² = 0.58, p < 0.0001) (Figure 9C), and negatively correlated with body weight (R² = 0.46, p = 0.0003) (Figure 9E). Additionally, 4-HNE levels exhibited a inverse correlation with physical activity (R² = 0.41, p = 0.0009) (Figure 9I). In contrast, MDA expression showed no meaningful associations with any of the five assessed physical phenotypes (R² < 0.4) (Figure 9K–O).

**Figure 9.**
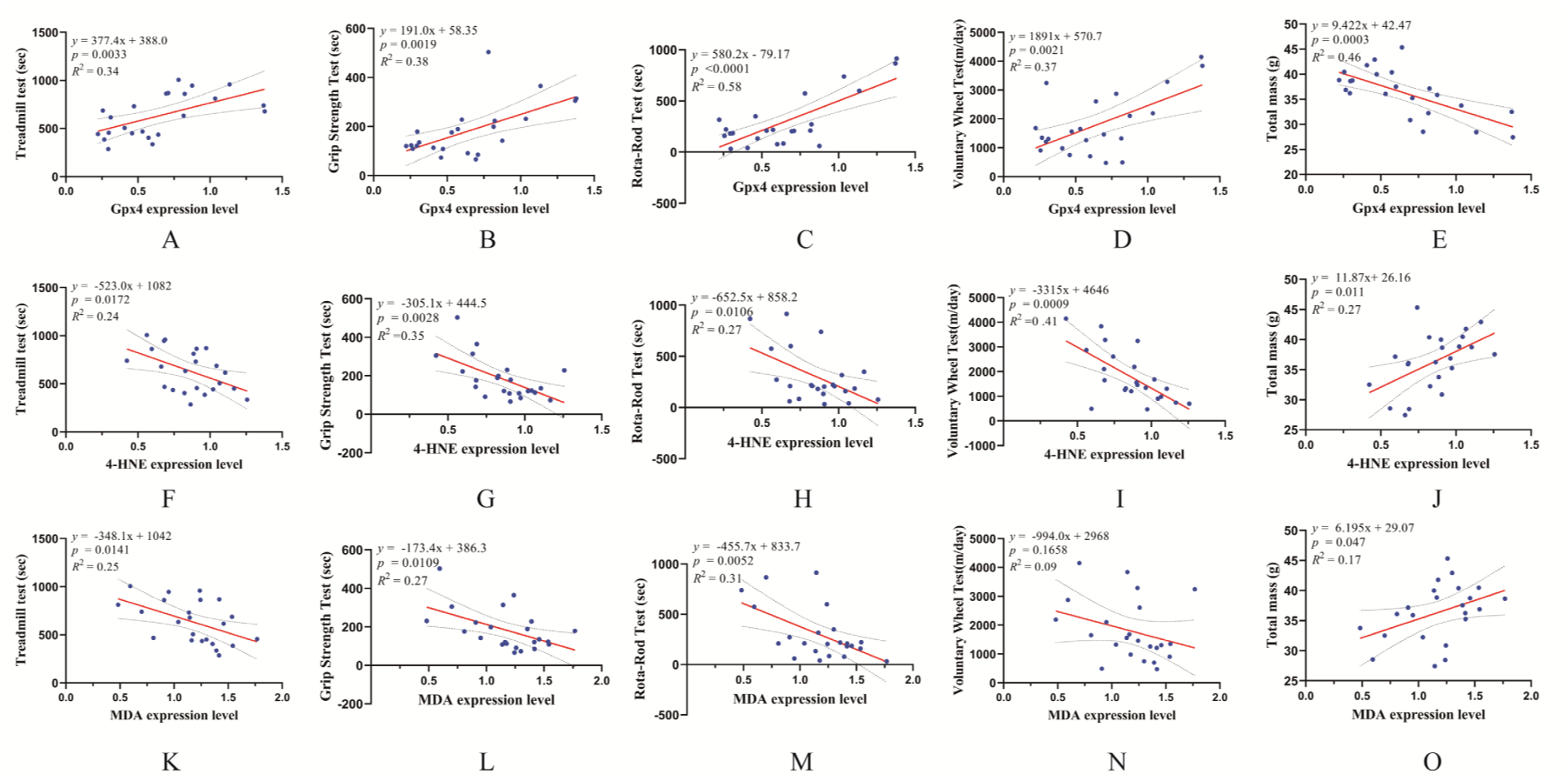
(A–E) Correlation analysis between GPX4 expression and endurance, grip strength, walking speed, physical activity, and body weight in total male mice. (F–J) Correlation analysis between 4-HNE expression and endurance, grip strength, walking speed, physical activity, and body weight in total male mice. (K–O) Correlation analysis between MDA expression and endurance, grip strength, walking speed, physical activity, and body weight in total male mice. R² values were calculated using Pearson correlation analysis, with values ≥ 0.4 considered indicative of a correlation and values ≥ 0.5 indicative of a strong correlation.

Compared with males, female mice showed distinct and overlapping correlation patterns between ferroptosis-related proteins and skeletal muscle physical phenotypes. Specifically, the negative correlation between GPX4 expression and body weight was more pronounced in females (R² = 0.63, p < 0.0001) (Figure 10E). Moreover, GPX4 expression in females was positively correlated with both endurance (R² = 0.58, p < 0.0001) and grip strength (R² = 0.54, p < 0.0001), relationships not observed in males (Figure 10A and B). In contrast to males, no correlation between GPX4 expression and walking speed was found in females (Figure 10E). Likewise, the negative correlation between 4-HNE expression and physical activity seen in males was absent in females (R² = 0.17, p = 0.473) (Figure 10I). Finally, only in females did MDA expression show correlations with endurance (R² = 0.63, p < 0.0001), grip strength (R² = 0.63, p < 0.0001), and body weight (R² = 0.63, p < 0.0001) (Figure 10K, L, and O).

**Figure 10.**
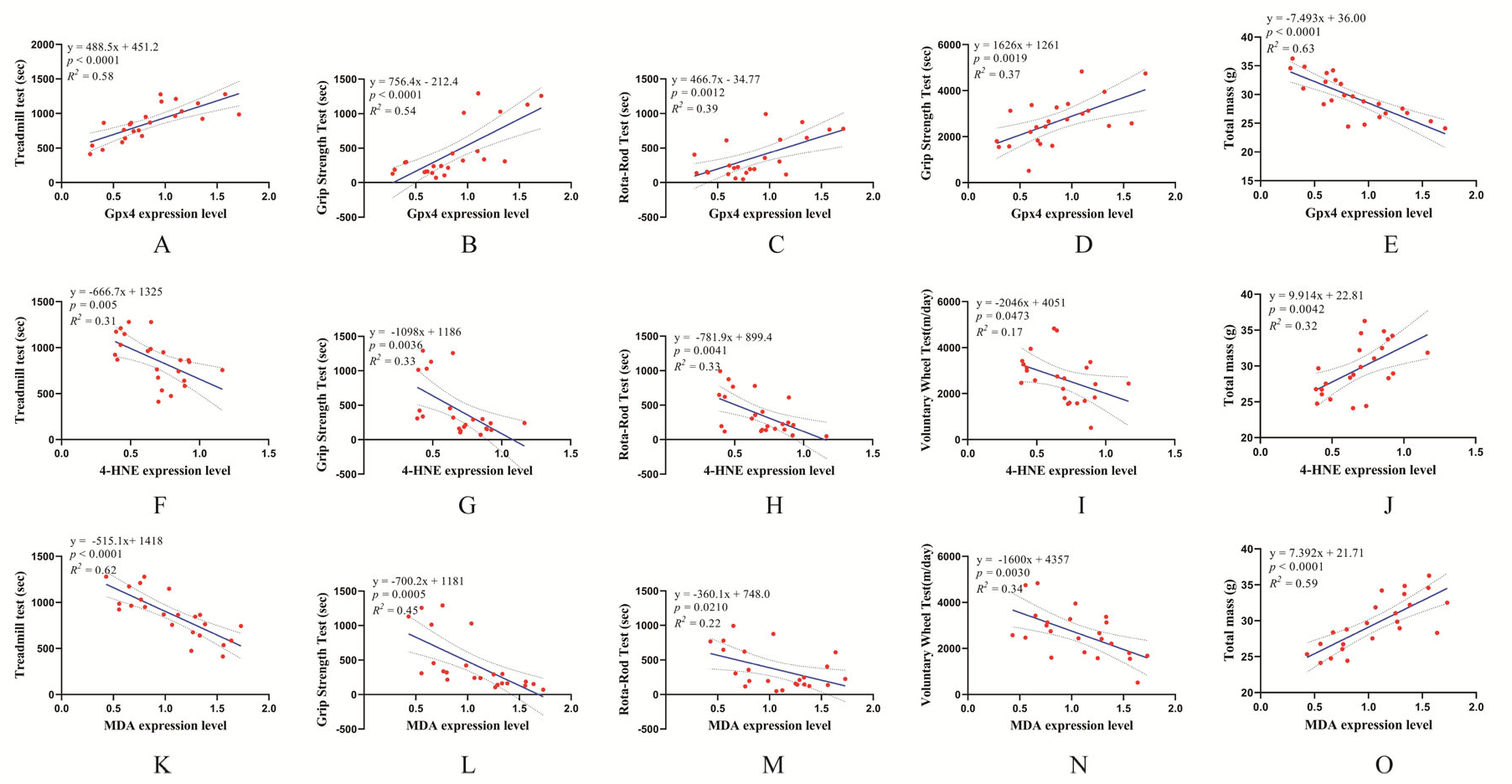
(A–E) Correlation analysis between GPX4 expression and endurance, grip strength, walking speed, physical activity, and body weight in total female mice. (F–J) Correlation analysis between 4-HNE expression and endurance, grip strength, walking speed, physical activity, and body weight in total female mice. (K–O) Correlation analysis between MDA expression and endurance, grip strength, walking speed, physical activity, and body weight in total female mice. R ² values were calculated using Pearson correlation analysis, with values ≥ 0.4 considered indicative of a correlation and values ≥ 0.5 indicative of a strong correlation.

These findings collectively indicate that ferroptosis-related signals are linked to multiple aspects of physical performance, with distinct and partially overlapping correlation patterns in males and females. Notably, GPX4 appears to serve as a central node linking ferroptosis defense to locomotor and strength phenotypes, particularly in aged female mice.

## Discussion

Ferroptosis is a regulated form of cell death influenced by a range of endogenous and exogenous stimuli, including aging and exercise [3, 18, 36–38]. However, the sex-specific regulation of ferroptosis-related markers in skeletal muscle during aging, as well as the differential effects of exercise modalities in aged populations, remain poorly understood. In this study, we first characterized sex-dependent differences in the regulation of ferroptosis-related markers across the aging process. We then examined the interaction between exercise modality and sex in modulating those markers in aged conditions. Our findings reveal that several ferroptosis-related markers are co-regulated by aging and sex, and in aged mice, some markers also display combined modulation by exercise modality and sex. These insights enhance the mechanistic understanding of sex-based variability in ferroptosis susceptibility and support the development of sex-specific personalized exercise interventions for aging-associated diseases.

In this study, mice were euthanized at 7 and 17 months of age, corresponding to human ages of 20–30 and 50–60 years, respectively [39, 40], to represent adulthood and the pre-senescent stage. These two age groups not only reflect physiological differences associated with aging but also provide an adequate time window for exploring future interventions in ferroptosis-related diseases. In terms of physical phenotypes, aging was accompanied by increased body weight, mainly attributable to higher fat mass and fat percentage. Concurrently, there was a marked decline in grip strength, walking speed, endurance, and overall physical activity, which further confirmed the aged phenotype across groups [23]. Notably, forced exercise produced more substantial improvements in specific physical phenotypes than voluntary exercise, with sex-specific effects evident in tissue fat mass, grip strength, and voluntary wheel activity.

Histological analyses using five distinct histological staining techniques revealed that four ferroptosis-associated markers—muscle fibrosis, 4-HNE and MDA accumulation, and Fe³⁺ deposition—were significantly elevated with aging. Masson’s trichrome staining showed a marked increase in collagen fibers in aged mice, indicative of skeletal muscle fibrosis [41]. It is important to note that several of the histological markers assessed in this study, such as fibrosis and 4-HNE accumulation, are not specific to ferroptosis and can also reflect broader processes, including chronic lipid peroxidation and proteostasis collapse. Therefore, our findings are better interpreted as indicating a “ferroptosis-related signature” and increased ferroptosis susceptibility, rather than definitive evidence that ferroptotic cell death is fully executed in aging skeletal muscle. Prior studies have implicated intracellular damage signals such as high-mobility group protein B1 in promoting fibrosis during ferroptosis [42–44]. Consistently, our IHC data show elevated 4-HNE and MDA, markers of oxidative stress and lipid peroxidation, in aged skeletal muscle [3, 45]. Prussian blue staining further confirmed excessive Fe³⁺ accumulation, reflecting impaired iron homeostasis and a potentiated Fenton reaction [46]. The Fenton reaction facilitates the formation of hydroxyl radicals through iron ion–catalyzed hydrogen peroxide decomposition, which triggers intracellular lipid peroxidation and consequently drives ferroptosis [47]. Collectively, these findings demonstrate that aging is a potent inducer of ferroptosis histopathology, with no overt sex differences at the tissue level.

Our exercise interventions revealed that forced exercise was significantly more effective than voluntary exercise in improving histological ferroptosis markers. Specifically, forced exercise led to a marked reduction in collagen fiber accumulation and oxidative damage irrespective of sex. However, Fe³⁺ levels exhibited a sex-specific response, with reductions confined to aged females undergoing forced exercise. These findings suggest that iron regulation, unlike other histological indicators, might be selectively modulated by the interaction between exercise modality and sex. In other words, aged females might experience superior improvements in iron homeostasis following forced exercise, potentially mitigating the risk of ferroptosis more effectively than in males in the same exercise condition.

Our protein-level analyses via WB and biochemical assays corroborated the histological findings. Aging reduced the expression of antioxidant and metabolic regulators such as GPX4, SLC7A11, and p-AMPK, while increasing oxidative damage indicators such as 4-HNE and MDA. These changes signify impaired antioxidant defenses and exacerbated lipid peroxidation, thereby predisposing aged muscle to ferroptosis. The absence of a detectable change in NCOA4 protein expression suggests that ferritinophagy may not be the predominant mechanism driving iron dysregulation in this model. Instead, our data point toward a scenario in which impaired GPX4–GSH redox defense and progressive iron accumulation create a pro-ferroptotic environment, without necessarily engaging all components of the canonical ferritinophagy pathway. Although most protein alterations were not sex-specific, GSSG and MDA levels were lower in aged females than aged males, indicating potentially superior redox homeostasis in females during aging [32].

Exercise-specific modulation of ferroptosis-related proteins was also evident. Forced exercise significantly altered the expression of GPX4, p-AMPK, and 4-HNE, with notable sex interactions. Aged female mice undergoing forced exercise showed higher GPX4 and lower 4-HNE than their male counterparts, suggesting enhanced antioxidant and anti–lipid peroxidation capacity. Although p-AMPK was elevated in both sexes following forced exercise, the p-AMPK/AMPK ratio was higher in females, implying a more favorable adaptation in lipid metabolism. Further biochemical analyses confirmed that Fe³⁺, GSSG, and MDA levels, as well as the GSH/GSSG ratio, were significantly influenced by the interaction between exercise modality and sex. Aged females demonstrated lower oxidative stress and improved redox balance than males under forced exercise, reinforcing the concept of sex-specific advantages in exercise responsiveness.

At the gene level, we analyzed 18 ferroptosis-related transcripts from antioxidant, iron, and lipid metabolism pathways. Among them, SCL7A11, GSS, GCLC, and POR exhibited significant alterations under the combined influence of aging and sex. Aging selectively downregulated SCL7A11 and GSS in females and GCLC in males, indicating divergent vulnerabilities in antioxidant pathways. POR, a pro-oxidative enzyme essential for ferroptosis [48], showed sex-specific expression despite being unaltered by aging, suggesting that it mediates sex differences in ferroptosis susceptibility. These findings indicate that certain ferroptosis-related gene markers are subject to sex-specific regulation during the aging process.

Post-exercise gene expression analyses revealed that 11 of the 18 genes were significantly regulated by the interaction between exercise modality and sex. In particular, SCL7A11, GSS, GCLC, SLC40A1, SCL11A2, POR, ACSL4, and LPCAT3 were co-regulated by both exercise modality and sex. Notably, forced exercise induced five distinct gene expression changes exclusively in aged females: increased expression of SCL7A11, GSS, and SLC40A1 and decreased expression of SCL11A2 and ACSL4. In contrast, GCLC was upregulated in aged males. Although both sexes showed altered POR and LPCAT3 expression, the magnitude of response was greater in females. These results underscore that aged female show a more robust transcriptional response to forced exercise than males, particularly in genes linked to antioxidant defense, iron transport, and lipid metabolism.

Across histological, protein, and transcriptional levels, our findings indicate that aging exerts a stronger influence on ferroptosis regulation in male skeletal muscle than in female. Moreover, forced treadmill exercise was more effective than voluntary wheel running in modulating ferroptosis, with females demonstrating greater responsiveness than males. Although cumulative running distance over the 8-week intervention did not differ significantly between voluntary wheel running and forced HIIT treadmill exercise, the higher and more structured intensity of HIIT likely underlies the more pronounced anti-ferroptotic effects observed in the forced exercise groups. These observations highlight both sex- and intervention-specific modulation of ferroptosis, prompting further exploration of the underlying mechanisms.

Mechanistically, our findings are consistent with dysregulation of the NRF2–GPX4–GSH antioxidant axis, which represents a central defense system against lipid peroxidation and ferroptosis. Aging in our study was accompanied by coordinated reductions in GPX4, SLC7A11, GSS, and GCLC at the protein and/or mRNA level, together with lower GSH/GSSG ratio and higher MDA, indicating an impaired glutathione-dependent redox capacity. Conversely, forced exercise, particularly in aged females, partially restored GPX4 and GSH-related markers, suggesting re-engagement of this ferroptosis-protective pathway.

The relative protection observed in aged females at both baseline and after forced exercise may further reflect sex hormone–related regulation of ferroptosis susceptibility. Estrogen has been reported to enhance antioxidant enzyme expression, modulate iron handling (e.g., ferroportin and ferritin regulation), and improve mitochondrial ROS buffering [12, 13]. These mechanisms are compatible with our observations of lower MDA and GSSG levels and more favorable regulation of GSH- and lipid metabolism–related genes in aged females compared with males. While we did not directly assess NRF2 activation, circulating sex hormones, or mitochondrial ROS in this study, our data support the hypothesis that sex-specific modulation of the NRF2–GPX4–GSH axis and iron metabolism contributes to the differential ferroptosis vulnerability between males and females during aging.

One major contributor to the sex differences in ferroptosis susceptibility may lie in the divergent hormonal milieu between males and females. Estrogen, in particular, has been shown to confer protective effects on skeletal muscle through various pathways, including the upregulation of antioxidant enzymes (e.g., superoxide dismutase, glutathione peroxidase), enhancement of mitochondrial efficiency, and modulation of iron metabolism via suppression of ferroportin degradation [12, 13, 49]. The decline in estrogen after menopause in humans [50, 51], as well as in animal models following ovariectomy [52, 53], has been associated with increased oxidative stress and iron accumulation. In our study, aged females retained higher antioxidant capacity, which may reflect either residual ovarian hormone activity at 17 months or compensatory adaptations in antioxidant signaling pathways that sustain redox homeostasis.

In addition to hormonal influences, intrinsic properties of skeletal muscle fibers contribute to sex-specific ferroptosis responses. Female skeletal muscle generally contains a greater proportion of oxidative type I fibers [54], which are more resistant to ROS-induced damage [55, 56]. In contrast, males typically possess a higher abundance of glycolytic type II fibers [54, 57], which are more susceptible to lipid peroxidation and mitochondrial dysfunction [55, 56]. These phenotypic differences may underlie the disparities in ferroptotic stress observed in our study. Supporting this notion, our previous findings demonstrated that white muscle fibers exhibit lower GPX4 and higher 4-HNE expression compared to red muscle fibers [8], suggesting a fiber-type-dependent vulnerability to ferroptosis.

Sex-specific differences in adiposity may further exacerbate susceptibility, especially in aging males. Greater fat deposition in aging males can promote inflammatory cell infiltration, fibrosis, and ROS accumulation—key triggers of ferroptotic signaling [8]. Thus, the convergence of hormonal, metabolic, and structural factors help explain the more pronounced ferroptotic alterations in aging male skeletal muscle.

Turning to the role of exercise interventions, forced treadmill exercise exerted stronger anti-ferroptotic effect than voluntary wheel running. Mechanistically, this superiority may be attributed to enhanced antioxidant defenses, improve lipid metabolism, and more efficient iron regulation [58, 59]. High-intensity endurance exercise has been reported to elevate fatty acid oxidation, reduce mitochondrial ROS emission, and lower lipid peroxidation susceptibility—all of which contribute to ferroptosis resistance. Forced exercise may also facilitate iron export and sequestration, thereby reducing labile iron pools, particularly in females [60].

These findings suggest that the protective benefits of forced HIIT against ferroptosis are amplified in females, likely owing to more efficient iron handling through the glutathione pathway, enhanced antioxidant adaptation, and greater transcriptional responsiveness. By contrast, aged males may require additional strategies—such as dietary iron modulation, antioxidant supplementation, or pharmacological ferroptosis inhibitors—to attain comparable protective effects.

Translationally, the more pronounced benefits of forced HIIT treadmill exercise compared with voluntary wheel running suggest that structured, externally guided exercise may be particularly effective in attenuating ferroptosis-related alterations in aging skeletal muscle. In this context, forced exercise in mice may parallel professionally supervised exercise interventions in humans, whereas voluntary wheel running more closely resembles self-guided activity. Notably, because cumulative running distance did not differ significantly between modalities, these differences are likely driven by training intensity and structure rather than volume alone, supporting the need for future clinical studies comparing supervised versus self-guided exercise programs in older or at-risk populations.

Our correlation analyses maybe support a functional relevance of ferroptosis signaling to physical phenotype in a sex-dependent manner. In male mice, higher GPX4 expression was associated with faster walking speed and lower body weight, whereas increased 4-HNE was linked to reduced spontaneous physical activity, suggesting that impaired antioxidant defense and enhanced lipid peroxidation may contribute to locomotor slowing and decreased habitual activity. By contrast, in females, GPX4 showed a broader pattern of favorable associations, being positively correlated not only with endurance but also with grip strength, while its inverse relationship with body weight was even stronger than in males. Moreover, only in females did MDA exhibit robust correlations with endurance, strength, and body weight, whereas no meaningful associations were observed in males. These findings suggest that, particularly in aged females, GPX4 and lipid peroxidation markers (MDA, 4-HNE) act as key molecular nodes coupling ferroptosis susceptibility to multiple domains of physical phenotypes, including locomotion, muscular strength, and body composition. Taken together, these sex-specific correlation patterns maybe reinforce the notion that ferroptosis-related redox imbalance is not merely a molecular hallmark of muscle, but is also tightly linked to clinically relevant phenotypes such as mobility and muscle strength, with females showing a closer coupling between ferroptosis defense and functional capacity than males.

A limitation of the present study is that exercise dose was not rigorously matched using physiological indices such as energy expenditure or VO₂-based measures. Although we quantified running distance in both voluntary wheel and forced HIIT treadmill groups and found no significant difference in cumulative distance over the 8-week intervention, we did not directly assess parameters such as %VO₂peak, %VO₂max, or total work. Therefore, the more pronounced anti-ferroptotic and phenotypic adaptations induced by forced exercise may, at least in part, be attributable to the distinct intensity profiles and the highly structured nature of the training protocol. Future studies incorporating precise monitoring of exercise intensity and energy expenditure will be necessary to disentangle the relative contributions of exercise dose versus exercise modality to ferroptosis regulation in aging skeletal muscle.

In conclusion, aging induces significant alterations in ferroptosis-related markers at the histological, protein, and gene levels, with notable sex-specific differences. Under aged conditions, forced exercise is more effective than voluntary exercise in mitigating ferroptosis risk, particularly in aged females. Enhanced antioxidant capacity, improved lipid metabolic adaptation, and better iron regulation underlie the superior responses observed in females. These findings provide a comprehensive mechanistic framework for sex-tailored exercise interventions aimed at mitigating ferroptosis and its associated pathologies in the aging population.

## Ethical approval and consent to participate

All the procedures followed in this experiment were approved by the Institutional Animal Care and Use Committee of Hanyang University (HYU 2021-0239A).

## Consent for publication

Not applicable.

## Availability of Data and Materials

The data used to support the findings of this study are presented here. Any further data needed are available from the corresponding author upon request.

## Declaration

The above data and conclusions are derived from a part of Fujue Ji’s doctoral dissertation.

## Author contributions

Fujue Ji: Conceptualization, data curation, formal analysis, investigation, methodology, writing—original draft.

Hyeonseung Rheem: Data curation and methodology.

Haesung Lee: Formal analysis and methodology.

Minyeong Eom: Visualization.

Jong-Hee Kim: Conceptualization, supervision, funding acquisition, writing—review and editing.

## Acknowledgments

We would like to thank Prof. Dr. Gwang-woong Go (Hanyang University, Korea) for generously providing the DEXA machine used for the body composition analysis of the mice and the PowerUp™ SYBR™ Green PCR master mix (Thermo Scientific, #A25742) for the RT-qPCR.

We are grateful for support in Skill Learning from Kaixin Doctor and MASCU (Medical Association with Science, Creativity, and Unity), Inc., Shenzhen, China.

## References

1. Tang, D. and G. Kroemer, Ferroptosis. Current Biology, 2020. 30(21): p. R1292–R1297.

2. Li, J., et al., Ferroptosis: past, present and future. Cell death & disease, 2020. 11(2): p. 88.

3. Tang, D., et al., Ferroptosis: molecular mechanisms and health implications. Cell Res, 2021. 31(2): p. 107–125.

4. Bebber, C.M., et al., Ferroptosis in Cancer Cell Biology. Cancers (Basel), 2020. 12(1).

5. Reichert, C.O., et al., Ferroptosis mechanisms involved in neurodegenerative diseases. International journal of molecular sciences, 2020. 21(22): p. 8765.

6. Huang, Y., et al., Ferroptosis in a sarcopenia model of senescence accelerated mouse prone 8 (SAMP8). International journal of biological sciences, 2021. 17(1): p. 151.

7. Sun, C.-C., et al., Ferroptosis and Its Potential Role in the Physiopathology of Skeletal Muscle Atrophy. International Journal of Molecular Sciences, 2024. 25(22): p. 12463.

8. Ji, F., et al., Differential ferroptosis regulation in red and white gastrocnemius under obesity and its Attenuation by exercise and dietary restriction. Scientific Reports, 2025. 15(1): p. 23821.

9. Finkel, T. and N.J. Holbrook, Oxidants, oxidative stress and the biology of ageing. nature, 2000. 408(6809): p. 239-247.

10. Alves, F.M., et al., Age-related changes in skeletal muscle iron homeostasis. The Journals of Gerontology: Series A, 2023. 78(1): p. 16–24.

11. Wei, Z., et al., Aging lens epithelium is susceptible to ferroptosis. Free Radical Biology and Medicine, 2021. 167: p. 94–108.

12. Ide, S., et al., Sex differences in resilience to ferroptosis underlie sexual dimorphism in kidney injury and repair. Cell reports, 2022. 41(6).

13. Liang, D., et al., Ferroptosis surveillance independent of GPX4 and differentially regulated by sex hormones. Cell, 2023. 186(13): p. 2748–2764. e22.

14. Ji, L.L., Exercise-induced modulation of antioxidant defense. Annals of the New York Academy of Sciences, 2002. 959(1): p. 82–92.

15. Souza, J., et al., Physical-exercise-induced antioxidant effects on the brain and skeletal muscle. Antioxidants, 2022. 11(5): p. 826.

16. Martinez-Huenchullan, S.F., et al., Skeletal muscle adiponectin induction in obesity and exercise. Metabolism-Clinical and Experimental, 2020. 102.

17. Ji, F., H.S. Lee, and J.-H. Kim, Resistance exercise and skeletal muscle: protein synthesis, degradation, and controversies. European Journal of Applied Physiology, 2025: p. 1–30.

18. Wang, P., et al., Acute exercise stress promotes Ref1/Nrf2 signalling and increases mitochondrial antioxidant activity in skeletal muscle. Experimental physiology, 2016. 101(3): p. 410–420.

19. Pellegrino, M.A., et al., Redox homeostasis, oxidative stress and disuse muscle atrophy. The Journal of physiology, 2011. 589(9): p. 2147–2160.

20. Alves, F.M., et al., Iron accumulation in skeletal muscles of old mice is associated with impaired regeneration after ischaemia–reperfusion damage. Journal of Cachexia, Sarcopenia and Muscle, 2021. 12(2): p. 476–492.

21. Scicchitano, B.M., et al., The physiopathologic role of oxidative stress in skeletal muscle. Mechanisms of Ageing and Development, 2018. 170: p. 37–44.

22. Ji, F. and J.-H. Kim, Muscle Type-Specific Modulation of Autophagy Signaling in Obesity: Effects of Caloric Restriction and Exercise. Journal of Obesity & Metabolic Syndrome, 2025. 34(3): p. 303.

23. Ji, F., et al., Overlapping and Distinct Physical and Biological Phenotypes in Pure Frailty and Obese Frailty. Bioscience Reports, 2024. 44(11).

24. Ji, F., et al., The impact of frailty syndrome on skeletal muscle histology: preventive effects of exercise. FEBS Open Bio, 2024.

25. Livak, K.J. and T.D. Schmittgen, Analysis of relative gene expression data using real-time quantitative PCR and the 2(-Delta Delta C(T)) Method. Methods, 2001. 25(4): p. 402–8.

26. Gupta, A., T.S. Stead, and L. Ganti, Determining a meaningful R-squared value in clinical medicine. Academic Medicine & Surgery, 2024.

27. Choueiry, G. What is a Good R-Squared Value? [Based on Real-World Data]. 2021 2021/11/13; Available from: https://quantifyinghealth.com/r-squared-study/.

28. Zhang, X., et al., High-fat diet alleviates colitis by inhibiting ferroptosis via solute carrier family seven member 11. The Journal of Nutritional Biochemistry, 2022. 109: p. 109106.

29. Zhao, X., et al., Adipose tissue macrophage-derived exosomes induce ferroptosis via glutathione synthesis inhibition by targeting SLC7A11 in obesity-induced cardiac injury. Free Radical Biology and Medicine, 2022. 182: p. 232–245.

30. Ding, H., et al., Transferrin receptor 1 ablation in satellite cells impedes skeletal muscle regeneration through activation of ferroptosis. J Cachexia Sarcopenia Muscle, 2021. 12(3): p. 746–768.

31. Mancardi, D., et al., Iron overload, oxidative stress, and ferroptosis in the failing heart and liver. Antioxidants, 2021. 10(12): p. 1864.

32. Giustarini, D., et al., Pitfalls in the analysis of the physiological antioxidant glutathione (GSH) and its disulfide (GSSG) in biological samples: An elephant in the room. Journal of Chromatography B, 2016. 1019: p. 21–28.

33. Tsikas, D., Assessment of lipid peroxidation by measuring malondialdehyde (MDA) and relatives in biological samples: Analytical and biological challenges. Analytical biochemistry, 2017. 524: p. 13–30.

34. Seibt, T.M., B. Proneth, and M. Conrad, Role of GPX4 in ferroptosis and its pharmacological implication. Free Radical Biology and Medicine, 2019. 133: p. 144–152.

35. Yang, W.S. and B.R. Stockwell, Ferroptosis: death by lipid peroxidation. Trends in cell biology, 2016. 26(3): p. 165–176.

36. Wang, H.-T., W.-Q. Yang, and Y.-Q. Liu, Effects of aerobic exercise on Nrf2/GPX4/ferroptosis pathway in myocardial injury in high-fat diet mice. Chinese Journal of Applied Physiology, 2022. 38(2): p. 143.

37. Espinoza, S.E., et al., Glutathione peroxidase enzyme activity in aging. The Journals of Gerontology Series A: Biological Sciences and Medical Sciences, 2008. 63(5): p. 505–509.

38. Mazhar, M., et al., Implication of ferroptosis in aging. Cell Death Discovery, 2021. 7(1): p. 149.

39. Justice, J.N., et al., Comparative Approaches to Understanding the Relation Between Aging and Physical Function. J Gerontol A Biol Sci Med Sci, 2016. 71(10): p. 1243–53.

40. Dutta, S. and P. Sengupta, Men and mice: relating their ages. Life sciences, 2016. 152: p. 244–248.

41. Mahdy, M.A., Skeletal muscle fibrosis: an overview. Cell and tissue research, 2019. 375(3): p. 575–588.

42. Yu, Y., et al., Hepatic transferrin plays a role in systemic iron homeostasis and liver ferroptosis. Blood, The Journal of the American Society of Hematology, 2020. 136(6): p. 726–739.

43. Zhao, X., et al., Adipose tissue macrophage-derived exosomes induce ferroptosis via glutathione synthesis inhibition by targeting SLC7A11 in obesity-induced cardiac injury. Free Radic Biol Med, 2022. 182: p. 232–245.

44. Zhang, X., et al., High-fat diet alleviates colitis by inhibiting ferroptosis via solute carrier family seven member 11. J Nutr Biochem, 2022. 109: p. 109106.

45. Chen, X., et al., Characteristics and biomarkers of ferroptosis. Frontiers in cell and developmental biology, 2021. 9: p. 637162.

46. Henle, E.S., Y. Luo, and S. Linn, Fe2+, Fe3+, and oxygen react with DNA-derived radicals formed during iron-mediated Fenton reactions. Biochemistry, 1996. 35(37): p. 12212–12219.

47. Shen, Z., et al., Fenton-reaction-acceleratable magnetic nanoparticles for ferroptosis therapy of orthotopic brain tumors. ACS nano, 2018. 12(11): p. 11355–11365.

48. Yan, B., et al., Membrane damage during ferroptosis is caused by oxidation of phospholipids catalyzed by the oxidoreductases POR and CYB5R1. Molecular cell, 2021. 81(2): p. 355–369. e10.

49. Liu, L., Y. Jiang, and S. Li, Research progress on the mechanism of sex hormones and their receptors in liver lipid metabolism. Chinese Journal of Endocrinology and Metabolism, 2020. 36: p. 267–272.

50. Cai, H., et al., Iron accumulation and its impact on osteoporotic fractures in postmenopausal women. Journal of Zhejiang University-Science B, 2023. 24(4): p. 301–311.

51. Ahanchi, N.S., et al., The complementary roles of iron and estrogen in menopausal differences in cardiometabolic outcomes. Clinical Nutrition, 2024. 43(5): p. 1136–1150.

52. Xiao, W., et al., Iron overload increases osteoclastogenesis and aggravates the effects of ovariectomy on bone mass. Journal of Endocrinology, 2015. 226(3): p. 121–134.

53. Liu, L.-l., et al., Iron accumulation deteriorated bone loss in estrogen-deficient rats. Journal of Orthopaedic Surgery and Research, 2021. 16(1): p. 525.

54. Nuzzo, J.L., Sex differences in skeletal muscle fiber types: A meta-analysis. Clinical anatomy, 2024. 37(1): p. 81–91.

55. Wang, F., et al., Effects of exercise - induced ROS on the pathophysiological functions of skeletal muscle. Oxidative medicine and cellular longevity, 2021. 2021(1): p. 3846122.

56. Powers, S.K., et al., Reactive oxygen species: impact on skeletal muscle. Comprehensive physiology, 2011. 1(2): p. 941–969.

57. Brooke, M.H. and W.K. Engel, The histographic analysis of human muscle biopsies with regard to fiber types: I. Adult male and female. Neurology, 1969. 19(3): p. 221–221.

58. Karlsson, J., Exercise, muscle metabolism and the antioxidant defense. World Review of Nutrition and Dietetics, 1997. 82: p. 81–100.

59. Noland, R.C., Exercise and regulation of lipid metabolism. Progress in molecular biology and translational science, 2015. 135: p. 39–74.

60. Buratti, P., et al., Recent advances in iron metabolism: relevance for health, exercise, and performance. Med Sci Sports Exerc, 2015. 47(8): p. 1596–1604.

